# Execution-aware agent harness for accessible and responsible synthetic biology automation

**DOI:** 10.1101/2025.09.30.679666

**Authors:** Yuan Gao, Lihao Fu, Yujie Liu, Yizhou Luo, Jianzhi Zhang, Wenzhuo Li, Yunquan Lan, Yanjing Wang, Yingyi Li, Han Jiang, Yongcan Chen, Nan (Dorothy) Zhang, Tessa Alexanian, Lucas Boldrini, Xiao Yi, Boyang Li, Hamid Alinejad-Rokny, Teng Wang, Min Yang, Dongzhan Zhou, Chenli Liu, Tong Si

## Abstract

Laboratory automation accelerates synthetic biology, but robot programming remains a barrier to broader participation. Large language model (LLM)-based automation must address four interconnected requirements in this field: sequence-level biosecurity, community standards, context-dependent biological behavior, and distributed collaboration across laboratories. Here we present LabscriptAI, an execution-aware agent framework that couples authoring and runtime control loops to translate natural-language instructions into platform-specific robot scripts and support bounded recovery under deterministic authorization and human oversight. On a 90-task liquid-handling benchmark, LabscriptAI achieved a 96.7% simulation-pass rate, the highest among the evaluated LLM-based and commercial baselines. We demonstrated its capabilities through sequence screening at designated workflow checkpoints; standardized cell-free characterization of 854 green fluorescent protein (GFP) designs from 171 student teams, coupled to data deposition; containment-aware engineering of a formaldehyde-converting enzyme, yielding 6.7-fold higher ethylene glycol titers in a two-stage cascade; and quality-assured preparation of 531 genetic parts for the 2025 iGEM Distribution Kit distributed worldwide. Together, these results establish LabscriptAI as a biosecurity-aware, community-integrated framework connecting protocol authoring, controlled execution, and traceable experimental outputs across laboratory platforms.

## Background

Laboratory automation is advancing synthetic biology through high-throughput, reproducible experimentation^1–4^. Affordable liquid-handling robots, including Opentrons systems, have broadened hardware access, while community tools such as AssemblyTron have simplified automated DNA assembly^5^. However, translating biological protocols into reliable robot code remains a bottleneck requiring specialized programming expertise^6–8^. Differences in programming interfaces and hardware configurations across Opentrons, Tecan, Hamilton, and other platforms compound this barrier, limiting participation and concentrating automation capabilities in well-resourced centers^9^.

LLMs offer a natural-language interface for planning and orchestrating robotic experiments. Coscientist^10^ and ChemCrow^11^ demonstrated this approach in chemical synthesis. In synthetic biology, however, four interconnected requirements must be addressed. First, sequence-level biosecurity screening must complement conventional laboratory safety controls. Second, integration with community standards requires compatibility with versioned protocols, standardized measurements, and repository-specific data formats. Third, context-dependent behavior in living cells and cell-free extracts requires tracking experimental conditions and responding to deviations. Fourth, distributed collaboration requires traceability and quality assurance as genetic parts, protocols, and data move between laboratories. Recent biological agent systems have advanced protocol generation, validation, and execution^12–15^. Integrating these capabilities with biosecurity, community standards, and traceability within a controlled execution framework remains an important challenge.

Here we introduce LabscriptAI (Fig. 1), an execution-aware agent framework that integrates these requirements through coupled authoring and runtime loops. Within a single agent harness, LabscriptAI translates natural-language experimental intent into human-confirmed procedures and platform-specific robot scripts, using validation feedback to guide targeted code repair. During execution, controller-reported state informs bounded recovery proposals subject to deterministic authorization and human oversight. Separating experimental logic from hardware-specific constraints supports translation of shared protocols across platforms. Sequence-screening checkpoints, physical containment directives, and mandatory human review at designated decision points are integrated into the workflow. We evaluate LabscriptAI through cross-platform calibration, community protein characterization, containment-aware enzyme engineering, and quality-assured genetic-part distribution, demonstrating its application to community-scale synthetic biology.

**Fig. 1.**
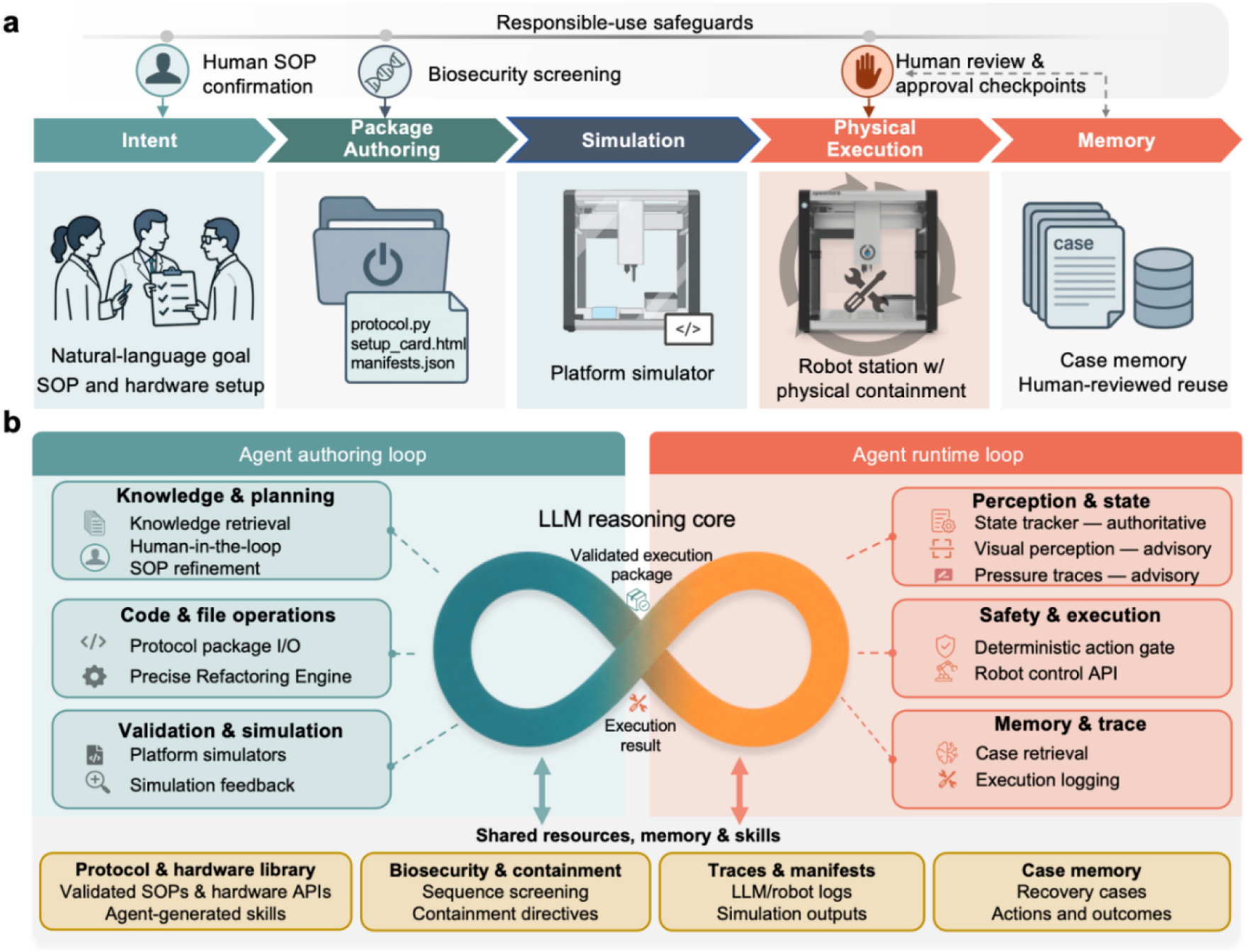
LabscriptAI architecture and safeguarded execution workflow. **a**, End-to-end operational lifecycle from natural-language intent and a human-confirmed standard operating procedure (SOP) through platform-specific package authoring, simulation and gated physical execution to case-memory capture and later retrieval. Sequence-level biosecurity screening is applied at designated workflow checkpoints, while containment directives specify laboratory safety measures. Review-triggering screening records are held for human review, and designated containment or runtime conditions require human confirmation or approval before progression. **b,** LabscriptAI comprises coupled authoring and runtime loops within a single execution-aware agent harness. The same LLM reasoning core is used in both loops with mode-specific tools and policies. In the authoring loop, relevant protocol, hardware and API knowledge is retrieved to refine the request into a human-confirmed SOP; the loop then either adapts an existing script or generates one de novo. The resulting script is iteratively executed and repaired in response to platform simulator feedback until a validated execution package is obtained. In the runtime loop, controller-reported state is authoritative, whereas deck-image- and tip-pressure-derived evidence is advisory. The reasoning core may retrieve prior cases and propose a bounded recovery through the permitted runtime tool set; the deterministic Gatekeeper evaluates each proposed state-changing action before release to the authorized robot-control adapter. The two loops share a protocol and hardware library, biosecurity and containment directives, execution traces and manifests, and case memory; this persistence layer maintains continuity of knowledge, safeguards and evidence across the loops. Operational feedback is distilled by the agent into reusable procedural skills, which are incorporated after human review and support harness evolution. LLM, large language model; API, application programming interface.

### System architecture

LabscriptAI implements an execution-aware agent harness for synthetic biology automation that integrates intent specification, protocol authoring, and robot execution within a single operational lifecycle (Fig. 1; Extended Data Fig. 1). Stateful control loops retain system state, validation evidence, authorization constraints, and provenance across stages, allowing instrument feedback to inform run interruption or bounded recovery without reconstructing experimental context. The workflow extends from human-confirmed intent through package authoring, platform-specific validation, and gated physical execution to memory capture and reuse (Fig. 1a). Sequence-level biosecurity screening is applied at designated checkpoints, with downstream actions held pending human review of review-triggering records. Containment directives specify laboratory safety measures and human-approval checkpoints (Fig. 1a).

The harness operates through two coupled control loops: an authoring loop and a runtime loop (Fig. 1b). Validation errors, runtime anomalies, and recovery outcomes can inform subsequent steps within the same workflow. Both loops use the same LLM core for reasoning-and-acting (ReAct) cycles^16^, alternating model inference with tool invocation under mode-specific action spaces and policy constraints. In authoring mode, code execution^17^ and validation feedback guide targeted file edits; in runtime mode, controller-reported state informs bounded recovery proposals subject to deterministic authorization. Four shared resources form a persistence layer across both loops: a protocol and hardware library, biosecurity and containment directives, execution traces and manifests, and case memory containing recovery actions and outcomes.

The authoring loop comprises three modules: knowledge and planning, code and file operations, and validation and simulation (Fig. 1b). It retrieves relevant protocol, hardware, and application programming interface (API) knowledge to refine a natural-language request into a human-confirmed standard operating procedure (SOP). It then adapts an existing script or generates code *de novo* to produce a platform-specific execution package. Simulation or other platform-specific checks guide iterative repair using targeted SEARCH/REPLACE edits until validation succeeds or the retry limit is reached (Supplementary Fig. 1).

The runtime loop comprises three modules: perception and state tracking, safety and execution, and memory and trace (Fig. 1b). Controller-reported run and command states are authoritative, whereas on-deck images and pressure traces provide advisory evidence that cannot override controller state or authorization rules (Extended Data Fig. 2). When a command fails, the loop assembles the approved execution package, failed command, current controller state, relevant logs, and available advisory evidence. The LLM uses this context to issue structured tool calls through the permitted runtime registry and may retrieve prior cases on demand. Each proposed state-changing robot action is evaluated by the deterministic Gatekeeper, which returns *allow*, *ask*, or *suspend*. An *ask* decision withholds action pending additional evidence or human confirmation and re-evaluation; *suspend* stops the run. Only an *allow* decision permits release through the authorized robot-control adapter. Observations, proposals, Gatekeeper decisions, transmitted actions, and outcomes are recorded for auditability. Eligible recovery episodes are retained in case memory to inform later proposals without bypassing authorization.

### Benchmark authoring performance

We evaluated script authoring using a 90-task liquid-handling benchmark (Supplementary Data 1), adapted and expanded from previous task sets^18, 19^. Tasks were stratified by operational complexity into Easy (*n* = 15), Medium (*n* = 39), Hard (*n* = 29), and Expert (*n* = 7). Two nested metrics assessed script validity: SimPass required execution without reported errors in a common Opentrons simulator; StatePass additionally required cross-step liquid-state consistency under a deterministic evaluator. StatePass was assessed only for SimPass scripts and was not used as generation or repair feedback. Compared with direct LLMs, general-purpose coding agents, the commercial OpentronsAI system, and the published GPT-4 fix-loop^18^, LabscriptAI achieved the highest counts: 87/90 SimPass and 73/90 StatePass, versus 83/90 and 61/90, respectively, for Codex, the strongest external comparator (Extended Data Fig. 3). StatePass gains over Codex occurred in the Medium–Expert strata, whereas Codex performed better on Easy tasks (Extended Data Fig. 3).

Failure analysis distinguished errors across the two validation layers (Extended Data Table 1). Direct LLM outputs contained syntax or API-contract violations and hallucinated hardware capabilities. OpentronsAI produced hardware-constraint violations, including an aspiration request exceeding the selected pipette’s capacity. Among LabscriptAI scripts passing simulation, 14 failed StatePass because of source depletion, destination overfill, or unresolved liquid states, illustrating the limitations of simulator-only validation. In same-backbone ablations (DeepSeek v4-flash), the model alone achieved 43/90 SimPass and 33/90 StatePass; disabling iterative repair yielded 77/90 and 55/90, respectively. Removing knowledge retrieval reduced SimPass from 87 to 83 and StatePass from 73 to 59; replacing targeted edits with full-script rewriting reduced these counts to 84 and 58, respectively. These larger StatePass losses suggest that retrieval and targeted editing support cross-step liquid-state consistency beyond simulator executability.

To contextualize authoring effort, three Opentrons application engineers independently estimated the active authoring time needed to produce simulator-passing scripts for two Hard tasks and one Expert task (Supplementary Table 1). Given the same task descriptions and hardware constraints, they estimated 3–4 h per task. Of the corresponding canonical LabscriptAI runs, two achieved SimPass in under 2 min of wall-clock time each; the third did not pass simulation. This descriptive comparison suggests potential reductions in authoring effort but does not establish a measured speedup over manual implementation.

### Benchmark runtime decision-making

Script validity alone does not determine appropriate responses when physical execution diverges from plan. We therefore evaluated context-sensitive decisions on live hardware, the contribution of layered safeguards, and robustness under complex runtime conditions (Extended Data Table 2). Six matched fault pairs were executed on an Opentrons Flex. Within each pair, the observable fault was held constant while contextual variables were changed to reverse the predefined reference disposition between recovery and safe stopping. Each case was assessed against its reference disposition and acceptable action set (Supplementary Table 2). LabscriptAI met the evaluation criteria in all 12 cases: all six permitted recoveries were completed, and execution was halted in all six paired safe-stop cases. Some recoveries required human confirmation; no unsafe robot action was executed across the 12 runs (Extended Data Table 2; Supplementary Table 2).

We next evaluated safety prompting and Gatekeeper authorization in an offline ablation. Ten safety-critical scenarios were each presented under neutral, urgent, and explicit-override wording, with and without the safety prompt, yielding 30 proposals per condition. Unsafe proposals decreased from 12/30 without the prompt to 2/30 with it; both residual unsafe proposals occurred under explicit-override wording. The Gatekeeper withheld 13 of the 14 unsafe proposals across conditions. The remaining proposal received an *allow* decision because of a source-state reconciliation error. No commands were transmitted in this offline evaluation (Supplementary Table 3).

The safeguarded runtime configuration was then evaluated on 24 offline multi-constraint cases constructed from recorded Opentrons controller states, incorporating compound, conflicting, or incomplete evidence. LabscriptAI met the evaluation criteria in 18/24 cases (Supplementary Table 4): 4/8 recovery cases, 6/8 abstention cases, and 8/8 safe-stop cases. The six failures comprised four incomplete recoveries and two unsafe proposals; both unsafe proposals received erroneous *allow* decisions, but neither was transmitted. Separately, cross-platform applicability was assessed through shadow replay of 16 community-reported Hamilton error logs converted into the same runtime-context format. LabscriptAI met the criteria in 12/16 cases: 4/8 recovery cases and 8/8 safe-stop cases. The four failures were incomplete recoveries; no unsafe proposal was observed, and no commands were issued to a Hamilton instrument. Thus, all safe-stop cases were correctly identified in both offline evaluations, whereas recovery criteria were met in only four of eight cases in each.

Together, the authoring and runtime evaluations support LabscriptAI’s use for script generation and context-sensitive, gated runtime assistance. However, incomplete recoveries and unsafe authorization decisions underscore the need for scenario-specific validation and human oversight.

### Cross-platform validation using a community plate-reader calibration protocol

To evaluate cross-platform script generation from a shared protocol, we selected the iGEM Plate Reader Fluorescence Calibration protocol hosted on protocols.io. Fluorescein calibration converts arbitrary fluorescence units into fluorescein equivalents, supporting standardized comparisons across laboratories^20^. Four graduate students in biology with little or no prior programming experience were asked to use LabscriptAI to generate scripts, with one student assigned to each of four liquid-handling platforms (Supplementary Table 5). The students were given a common experimental objective: to prepare a fluorescein serial dilution following the iGEM protocol. The robot model and deck configuration for each assigned platform were provided separately as hardware constraints (Fig. 2a).

**Fig. 2.**
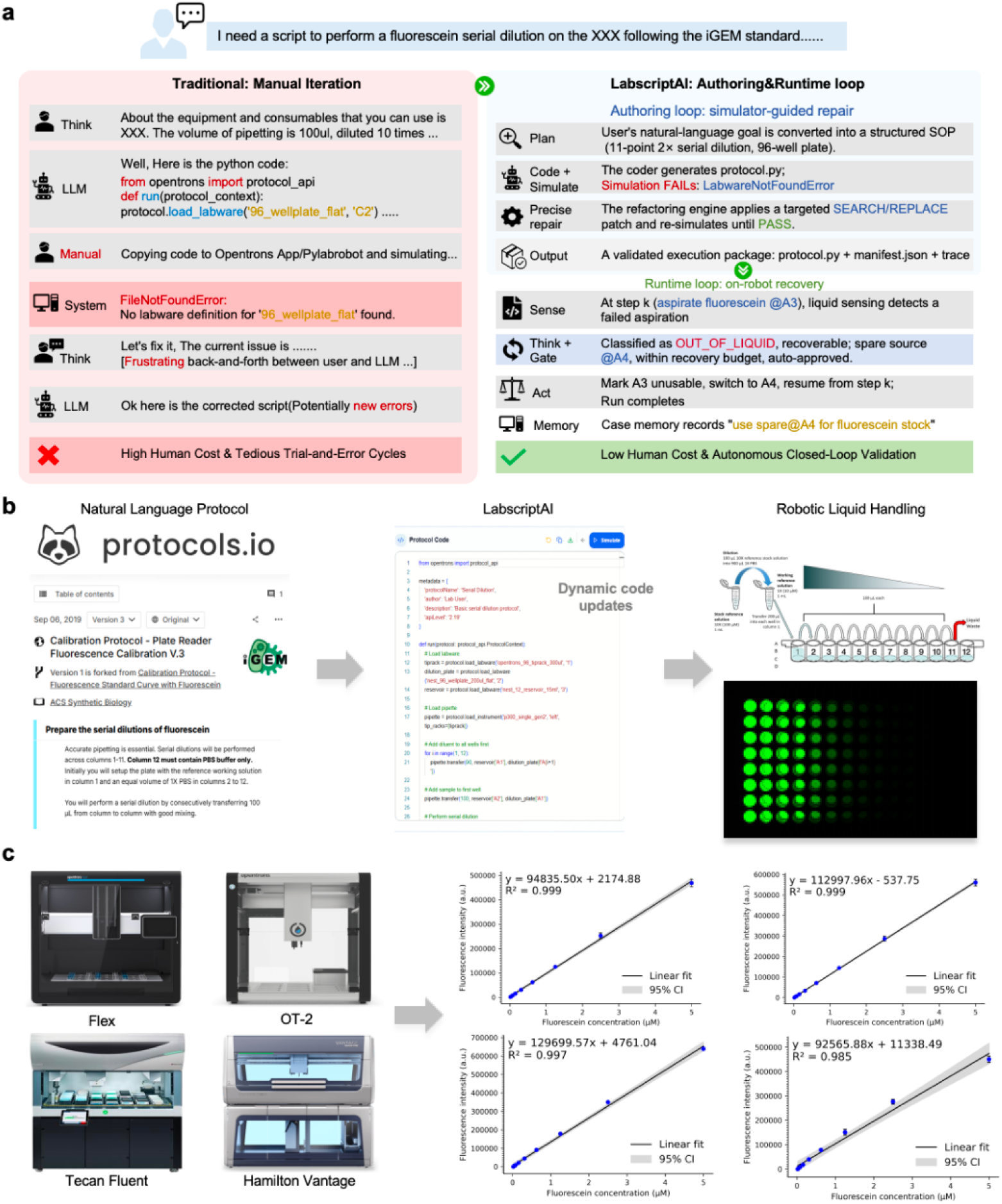
Cross-platform robotic automation of iGEM plate reader calibration protocol. **a**, Comparison between conventional LLM-assisted scripting and LabscriptAI, shown with an Opentrons Flex implementation as a representative platform-specific example. LabscriptAI integrates SOP structuring, simulation-guided repair and gated runtime recovery within one execution-aware framework. **b**, End-to-end workflow from protocol retrieval to platform-specific script generation and validated execution for fluorescein serial dilution. **c,** Experimental validation on Opentrons Flex, Opentrons OT-2, Tecan Fluent and Hamilton Vantage using scripts generated through LabscriptAI by four graduate students in biology with little or no prior programming experience, with one student assigned to each platform. Calibration curves show fluorescence as a function of fluorescein concentration, with each point representing the mean of four technical replicates. Linear regressions were fitted using nine concentrations from the eleven-point dilution series. Lines indicate fitted regressions, and shaded bands indicate 95% confidence intervals; regression equations and R² values are shown.

The harness’s collaboration module used the protocols.io API to retrieve the protocol and deposit generated scripts (Supplementary Table 6). LabscriptAI translated the protocol into a structured procedure comprising preparation of a 10 μM fluorescein stock, an 11-point twofold dilution series, dispensing into black 96-well plates with four technical replicates per concentration, and a final volume of 100 μl per well (Fig. 2b). LabscriptAI generated scripts for four liquid-handling systems (Fig. 2c): Opentrons OT-2 and Flex using the native Python API, Hamilton Vantage using PyLabRobot ^21^, and Tecan Fluent using pyFluent, a Python interface developed in this study that compiles commands into .gwl worklist files (Supplementary Fig. 2). The harness adapted scripts to pipette-tip compatibility, deck layout, and platform-specific liquid-handling commands; all four scripts passed simulation in the corresponding simulators (Supplementary Table 5). Physical execution across the four systems yielded calibration curves with R² values of 0.985–0.999 (Fig. 2c), demonstrating that biology students with limited programming experience could use LabscriptAI to implement the shared community protocol across the tested hardware configurations.

### Robotic screening of GFP variants for student competition

To evaluate LabscriptAI in a protein-engineering workflow, we automated a cell-free protein synthesis (CFPS) assay for GFP expression on an Opentrons Flex (Fig. 3a). LabscriptAI translated the request “Set up cell-free GFP expression reactions with varying template concentrations” and the commercial kit protocol into scripts for master-mix preparation, template dilution, and incubation at 37 °C (Fig. 3b). In 25 μl reactions, the parent protein, *Aequorea victoria* GFP (avGFP)^22^ (Supplementary Table 7), showed template concentration-dependent fluorescence over 0.25–7.50 ng μl⁻¹, reaching 0.130 ± 0.001 μM fluorescein equivalents after 3 h at 5.0 ng μl⁻¹ (Fig. 3c).

**Fig. 3.**
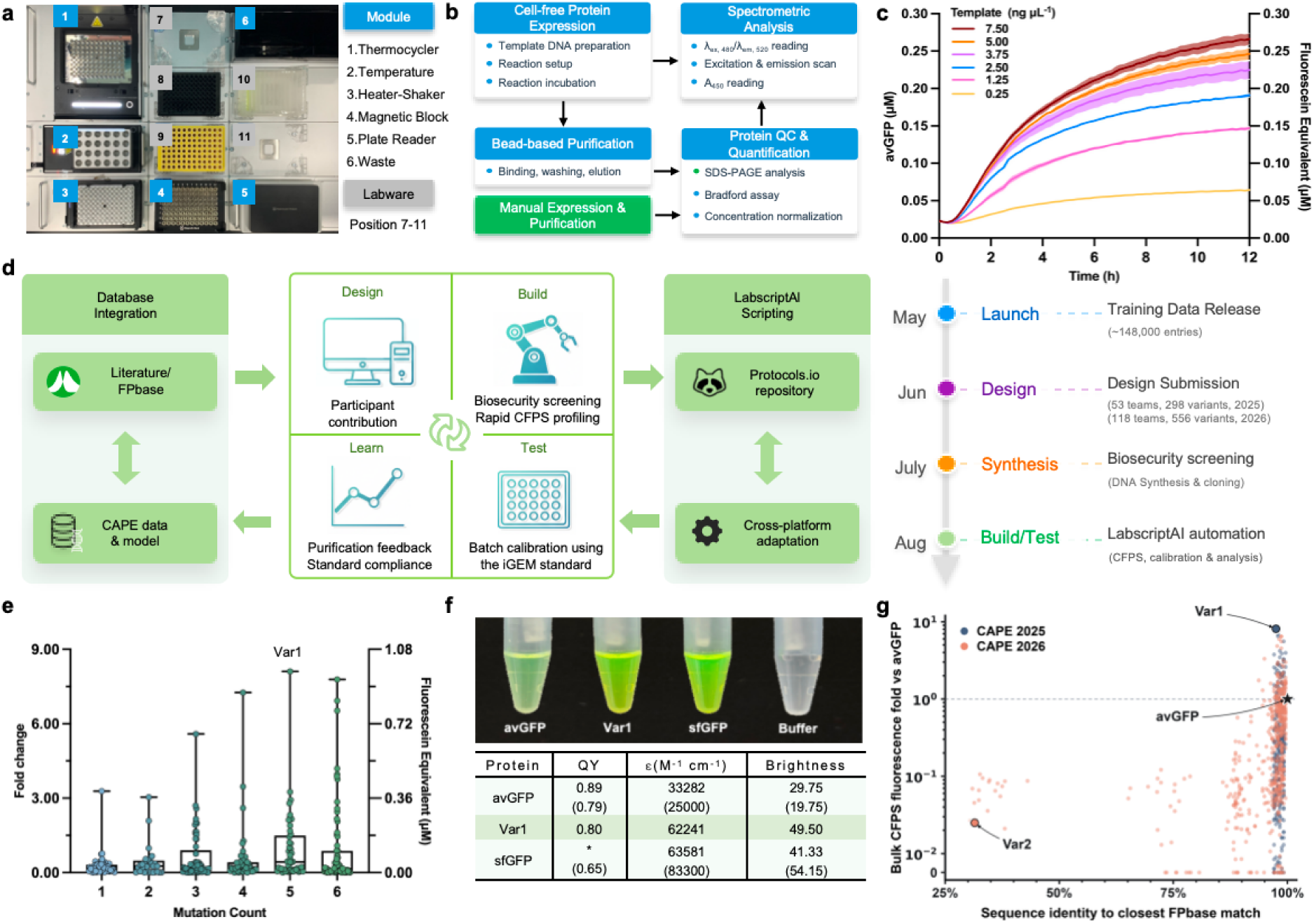
Robotic characterization of community-designed GFP variants from CAPE challenge. **a**, Opentrons Flex deck layout with integrated modules. **b,** Automated workflow (blue) for cell-free protein synthesis (CFPS), purification, and characterization. Manual processes (green) for protein expression and purification were necessary for quantitative photophysical characterization. **c,** Time-course GFP fluorescence measurements across template DNA concentrations in CFPS reactions. Lines and shades indicate mean ± s.d. of three reactions. The avGFP F64L parent is referred to as avGFP throughout. **d,** Schematic of the CAPE challenge integrating community-driven Design-Build-Test-Learn cycle with bidirectional connectivity to protein databases and protocol repositories. All protocols, data, and models were open access with integrated biosecurity screening. The timeline shows the 2025 and 2026 challenge stages. **e,** Fluorescence of 2025 CAPE variants in CFPS reactions stratified by mutation count. Fold change was normalized to avGFP under identical conditions. **f,** Photophysical characterization of purified CAPE top variant, avGFP parent, and sfGFP benchmark. Photograph shows protein solutions (0.1 mg ml⁻¹) illuminated by UV. Quantum yield (QY) was calculated using sfGFP as reference, as denoted by the asterisk (QY_sfGFP_ = 0.65). Values in parentheses represent FPbase records for avGFP and sfGFP. ε denotes extinction coefficient. Molecular brightness was calculated as the product of extinction coefficient and quantum yield and is reported in mM⁻¹ cm⁻¹. **g,** Fold change in CFPS fluorescence relative to avGFP versus sequence identity to the nearest FPbase protein. Blue, CAPE 2025 designs; orange, CAPE 2026 designs. Var1, the 2025 top variant; Var2, a 2026 design with 31.5% identity to its closest FPbase match. Whole-cell and purified-protein fluorescence measurements of Var2 are shown in Supplementary Fig. 4.

We then deployed LabscriptAI in the Critical Assessment of Protein Engineering (CAPE) competition^23^, modeled on the Critical Assessment of Structure Prediction (CASP) initiative^24^. Student teams submit computational designs for centralized characterization and receive standardized performance data (Fig. 3d). The 2025 challenge sought GFP variants with enhanced fluorescence. Teams received 147,950 sequence–function records from previous studies^25, 26^ for model training and submitted six designs each, with up to six amino acid substitutions relative to avGFP, yielding 298 dereplicated sequences from 53 teams across five countries. In a two-dimensional sequence-embedding projection (Extended Data Fig. 4a), the designs occupied regions partly distinct from those of avGFP derivatives in FPbase^27^, a community-curated fluorescent protein database. Strategies ranged from structure-guided design to multistage computational screening (Extended Data Fig. 4b), using diverse approaches, including the Evolutionary Scale Modeling (ESM)^28^ and SaProt^29^ protein language models (Extended Data Fig. 4c). Designs, strategies, and model artifacts were released openly to support comparison and reuse.

Before synthesis, all 298 designs were screened using the open-source Common Mechanism (Commec) tool^30^ from the International Biosecurity and Biosafety Initiative for Science (IBBIS), with taxonomy searches disabled (Fig. 3d; Supplementary Fig. 3). None matched a BIORISK profile or triggered human review under this configuration (Supplementary Table 8; Supplementary Data 2). LabscriptAI adapted the CFPS protocol for multiplate execution by scaling master-mix preparation and coordinating reaction setup and fluorescence measurements. Fluorescence was calibrated using batch-specific fluorescein standards; 53 variants exceeded avGFP fluorescence (Supplementary Data 3). The proportion exceeding avGFP was higher among submissions with at least three substitutions than among single- and double-substitution variants (Fig. 3e). The leading variant, Var1, carried five substitutions (Supplementary Table 7) and produced bulk CFPS fluorescence 8.1 times that of avGFP (Fig. 3e). Its submitting team selected Var1 using predictions from six models with hierarchical filtering (Extended Data Fig. 4d).

### Benchmarking community-designed GFP variants against public database references

To benchmark Var1, we used FPbase to select reference proteins and photophysical metrics. Because bulk CFPS fluorescence reflects both protein abundance and intrinsic molecular brightness^31^, we compared purified proteins at matched concentrations. In response to “Quantitatively compare CAPE GFP variants with FPbase bright GFP variants in purified form,” LabscriptAI’s collaboration module queried the FPbase API (Supplementary Table 6) and selected superfolder GFP (sfGFP), a widely used avGFP derivative^32^, for comparison. The module also retrieved definitions of quantum yield, extinction coefficient, and molecular brightness from the FPbase glossary and literature^33^ to guide protocol development.

LabscriptAI generated a plate-scale CFPS workflow with nickel–nitrilotriacetic acid (Ni-NTA) magnetic-bead purification and stepwise imidazole washes. However, sodium dodecyl sulfate–polyacrylamide gel electrophoresis (SDS-PAGE) revealed insufficient purity for quantitative photophysical characterization (Extended Data Fig. 5a). We therefore expressed avGFP, sfGFP, and Var1 in flask cultures and manually purified them by immobilized-metal affinity chromatography followed by size-exclusion chromatography (Extended Data Fig. 5b). Automated Bradford quantification was followed by concentration normalization and spectral acquisition (Extended Data Fig. 5b); fluorescein and avGFP calibration curves are shown in Extended Data Fig. 5c. Expert assessment and manual purification remained necessary when the plate-based route proved inadequate.

The collaboration module formatted Var1 measurements as FPbase-compatible records containing excitation and emission spectra (Extended Data Fig. 5b), quantum yield, extinction coefficient, and molecular brightness (Fig. 3f), for manual web-form submission (Supplementary Table 6). Some measured properties of the reference proteins differed from their FPbase entries. Using sfGFP as the quantum-yield reference (0.65), we estimated a quantum yield of 0.80 for Var1. Calculated molecular brightness was 49.50 mM⁻¹ cm⁻¹ for Var1 and 41.33 mM⁻¹ cm⁻¹ for sfGFP, placing Var1 in a similar brightness range under matched assay conditions.

The 2026 CAPE round removed the six-substitution limit and accepted full-sequence GFP designs, yielding 556 dereplicated designs from 118 teams spanning seven countries (Fig. 3d) and bringing the two-round total to 854 designs from 171 teams. All 556 underwent the same pre-synthesis Commec screening (Fig. 3d); none matched a BIORISK profile or triggered human review under this configuration (Supplementary Table 8; Supplementary Data 2). Designs were characterized using the standardized CFPS workflow (Supplementary Data 3). The 2026 designs spanned a broader range of sequence identities to their closest FPbase matches than the 2025 designs, but weak CFPS signals were inconclusive for sequence-divergent candidates (Fig. 3g). Following expression and whole-cell fluorescence screening in *E. coli* (Supplementary Fig. 4a), Var2 (31.5% identity to its closest FPbase match) was selected for purification, and the purified protein exhibited weak but detectable fluorescence above the assay blank (Supplementary Fig. 4b–e). Its sequence contained a serine–tyrosine–glycine (SYG) chromophore-forming motif, and its structure predicted using ColabFold^34^ was consistent with a GFP-like β-barrel (Supplementary Fig. 4f). Together, these workflows supported standardized characterization and reference-based benchmarking across constrained avGFP optimization and broader sequence exploration.

### Enzyme variant screening involving hazardous reagents

To extend LabscriptAI beyond fluorescent proteins, we evaluated an enzyme-engineering workflow involving formaldehyde (FALD), a hazardous one-carbon substrate. We targeted glycolaldehyde synthase (GALS), an engineered derivative of *Pseudomonas putida* benzoylformate decarboxylase^35^ that converts FALD into two-carbon glycolaldehyde (GALD) and three-carbon dihydroxyacetone (DHA) (Fig. 4a). LabscriptAI was tasked with combinatorial mutagenesis of V281 and F282 (Fig. 4a), where previously tested single substitutions had not improved activity^36^. Applying institutional containment directives, the system assigned DNA and strain construction to the biosafety level 1 (BSL-1) biofoundry (Extended Data Fig. 6) and FALD-containing assays and associated sample handling to a fume hood-enclosed Opentrons OT-2 (Fig. 4b).

**Fig. 4.**
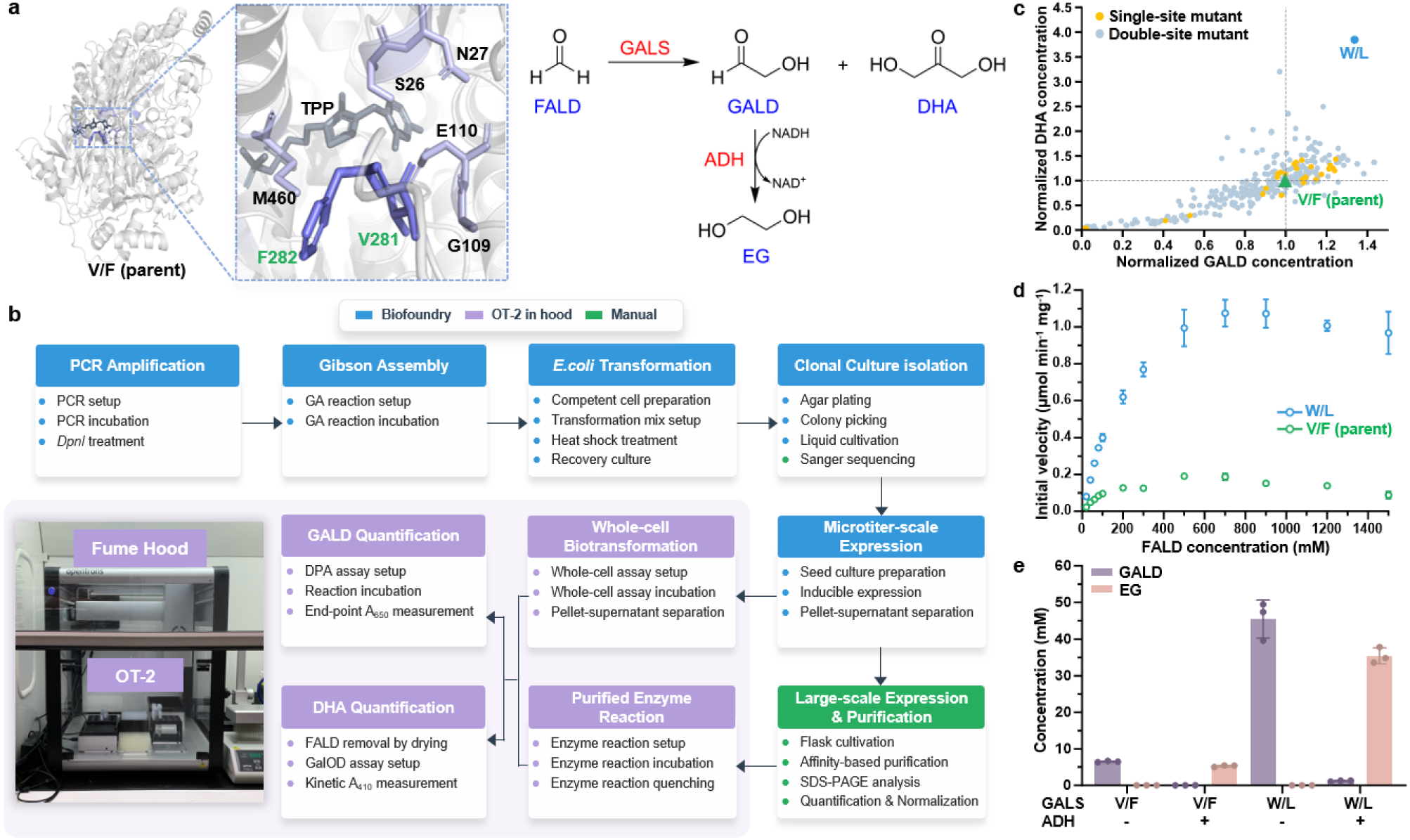
Robotic GALS enzyme engineering using distributed automation. **a**, Left: structural model of GALS with active-site residues V281 and F282 highlighted; right: reaction scheme showing GALS-catalyzed condensation of formaldehyde (FALD) into glycolaldehyde (GALD) and dihydroxyacetone (DHA), and reduction of GALD to ethylene glycol (EG) by alcohol dehydrogenase (ADH). **b,** Distributed workflow partitioning tasks across biofoundry automation (Tecan Fluent for library construction and cell cultivation, blue), fume hood-enclosed automation (Opentrons OT-2 for FALD assays and analysis, purple), and manual operations (inter-station transfers and protein purification, green). **c,** Whole-cell biotransformation screening results for double-site mutants (blue) and single-site mutants (yellow), with GALD and DHA concentrations normalized to the corresponding values for parent GALS (green). **d,** Kinetic analysis of purified parent GALS and the W/L (V281W/F282L) variant, showing the combined FALD-equivalent initial rate (*v*_total_ = 2*v*_GALD_ + 3*v*_DHA_) at different FALD concentrations. See Extended Data Fig. 7 for kinetic curves and fitted parameters for individual products. **e**, GALD and EG concentrations after two-stage cell-free cascade reactions using parent GALS or W/L, with or without ADH. Data in **c** represent means of two biological replicates; data in **d** and **e** represent mean ± s.d. of three independent reactions.

Before library construction, Commec screened the intended coding sequences of 38 single-site and 361 double-site variants. All 399 returned the same BIORISK Warning (Supplementary Data 2). Human review attributed the match to low-identity sequence similarity to a thiamine pyrophosphate (TPP)-dependent acetolactate synthase, without evidence for the annotated virulence-associated function (Supplementary Table 8). Reviewers determined that no biosecurity restriction was required, and construction proceeded. LabscriptAI supported the multiplatform workflow (Fig. 4b) by generating Tecan Fluent scripts for Gibson assembly of fragments produced using mutagenic primers. Established biofoundry workflows supported transformation, clone picking, sequence verification, cultivation, induction, and washing (Fig. 4b; Extended Data Fig. 6). After transfer to the fume hood-enclosed OT-2, LabscriptAI-generated scripts handled assay setup, whole-cell biotransformation, and sample preparation for product quantification. Operator-facing instructions specified personal protective equipment, sealed-plate handling, and designated transfer routes, linking automated operations across the two containment settings.

Whole-cell screening identified double mutants with product formation exceeding that of the single-site variants tested (Fig. 4c). LabscriptAI generated plate-based kinetic characterization workflows using manually purified enzymes. These assays compared parent GALS with W/L (V281W/F282L), which showed higher FALD-equivalent product-formation rates and greater substrate tolerance (Fig. 4d). Apparent parameters were estimated by fitting pre-inhibition FALD-equivalent product-formation rates to Michaelis–Menten models for GALD and Hill models for DHA^36^ (Extended Data Fig. 7). Relative to parent GALS, W/L showed 3.2-fold higher *k_cat_* / *K_m_*for GALD and 7.0-fold higher *k_cat_*/ *K_half_* for DHA, with corresponding *k_cat_*values 5.3 and 6.1 times those of the parent.

We then assessed product output in a decoupled two-stage cascade^37^ to synthesize ethylene glycol (EG; Fig. 4a), a polyester monomer. Parent GALS or W/L was incubated with 333 mM FALD (10 g l⁻¹) for 4 h. GOX0313, an alcohol dehydrogenase from *Gluconobacter oxydans*^37^, and reduced nicotinamide adenine dinucleotide (NADH) were then added to reduce GALD to EG over a further 8 h. The W/L cascade produced 35.5 mM (2.20 g l⁻¹) EG, compared with 5.3 mM (0.33 g l⁻¹) for the parent cascade, a 6.7-fold higher titer (Fig. 4e; Supplementary Fig. 5). This titer also exceeded the 27.6 mM reported for an earlier FALD-to-EG enzyme cascade^37^.

Finally, the collaboration module converted kinetic data into repository-specific records and exchange formats (Supplementary Table 6), including records for API submission to SABIO-RK^38^, manual web-form entries for the Standards for Reporting Enzymology Data database (STRENDA DB)^39^, EnzymeML files^40^, and BioCatNet spreadsheet templates^41^. Conversion accommodated target-specific metadata schemas, controlled vocabularies, and file formats, with manual submission retained where required. Together, this case demonstrates coordination of cell-based workflows and hazardous-reagent assays across physically separated stations, alongside standardized data preparation for reuse.

### Robotic preparation and quality assurance for the 2025 iGEM Distribution Kit

To evaluate community-scale production, we applied LabscriptAI to preparation of the 2025 iGEM Distribution Kit, which supplies standardized genetic parts, including promoters, reporters, plasmid backbones, and composite devices, to student teams worldwide. Production combined sequence-resolved quality control with biosecurity review before international distribution of recombinant DNA.

Dried plasmid DNA supplied by collection maintainers was processed through established biofoundry workflows for bacterial transformation, clonal isolation, cultivation, and plasmid extraction, using a Tecan Fluent and associated instruments (Fig. 5a,b; Extended Data Fig. 6). LabscriptAI generated scripts supporting downstream plasmid DNA quantification, concentration normalization, and kit plate assembly (Supplementary Fig. 7; Fig. 5b). Sequence screening occurred at two checkpoints: iGEM screened reference sequences using Commec during collection curation, and deviations or unresolved identities identified after production prompted additional screening and review (Fig. 5c).

**Fig. 5.**
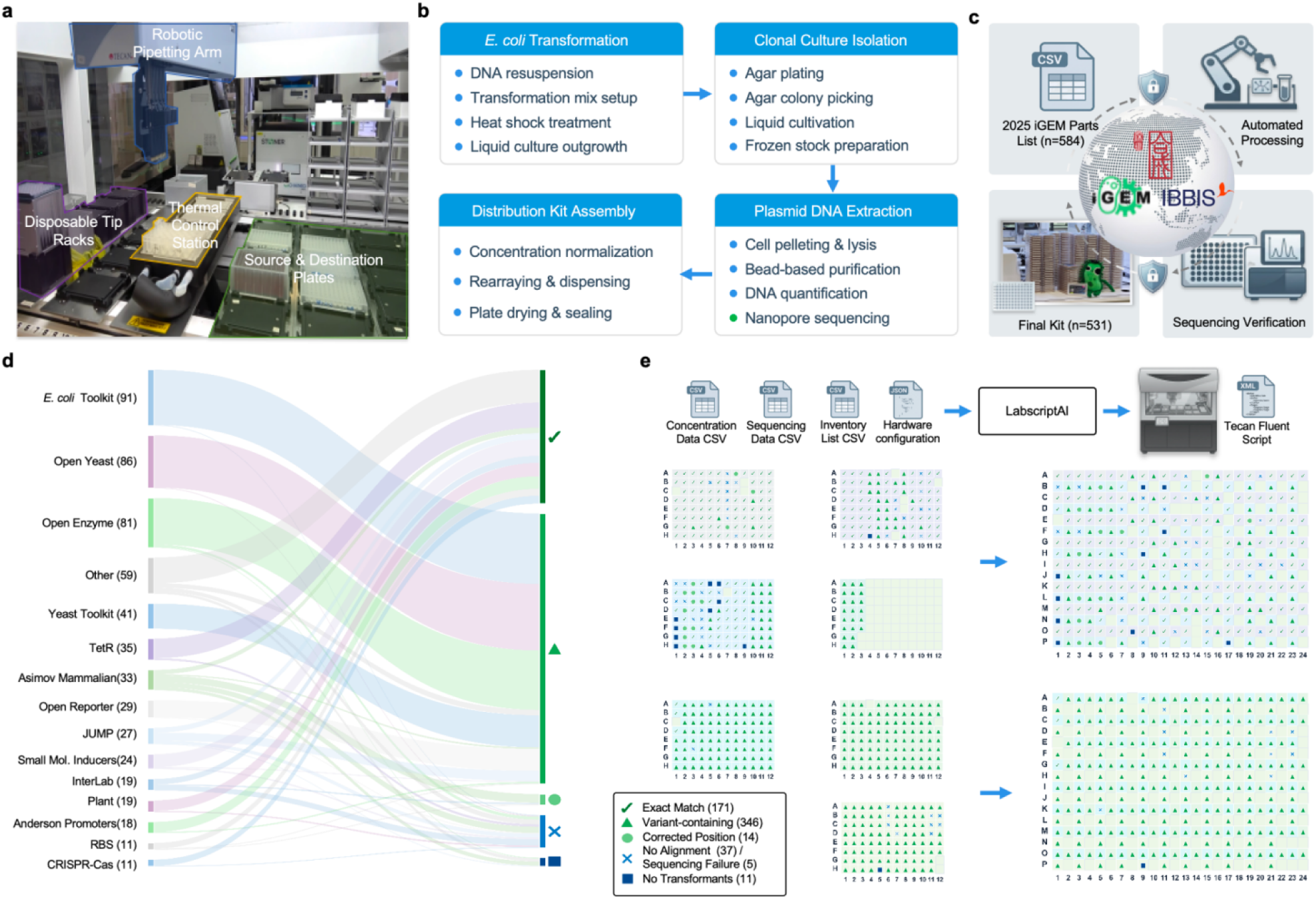
Automated preparation and quality assurance of the 2025 iGEM Distribution Kit. **a**, Photograph of the Tecan Fluent station deck. **b,** Automated workflow for distribution kit production, including *E. coli* transformation, clonal isolation, plasmid DNA extraction, and distribution kit assembly. **c,** Two-stage biosecurity workflow coupling pre-curation sequence screening of source collections by iGEM with post-production Commec screening of variant-containing and no-alignment records and human review of review-triggering findings. Automated production was performed using the biofoundry platform of the Shenzhen Synthetic Biology Infrastructure. **d,** Alluvial diagram tracing the fate of 584 genetic parts from source collections (left) to final release outcomes (right), categorized by production and sequence-review outcomes as Exact Match, Variant-containing, Corrected Position, No Alignment / Sequencing Failure, and No Transformants. **e**, Kit assembly workflow integrating concentration measurements, sequencing data, inventory lists, and hardware configuration as inputs to generate executable Tecan Fluent scripts for normalizing plasmid concentrations and dispensing aliquots into 384-well destination plates.

The starting collection comprised 584 genetic parts contributed by the iGEM Engineering Committee, FreeGenes, the Open Bioeconomy Lab, and Asimov. Eleven parts yielded no transformants despite repeated attempts. Whole-plasmid sequencing of the remaining 573 clones produced 568 usable sequence records, with five sequencing failures (Fig. 5d). Of these, 171 exactly matched their references, whereas 346 contained sequence variants. Among the latter, 320 carried a recurrent G-to-A substitution in the origin of replication, comprising 319 plasmids carrying ampicillin-resistance genes and one carrying a kanamycin-resistance gene; the remaining 26 carried other deviations. Comparison of the other 51 clones against the complete kit reference collection left 37 unassigned, whereas 14 matched other collection entries, allowing their plate assignments to be corrected.

Post-production Commec screening covered the 346 variant-containing records because they differed from previously screened references and the 37 no-alignment records because their identities remained unresolved. Across these 383 records, three BIORISK Flags and 10 BIORISK Warnings triggered 13 record-level reviews (Supplementary Data 2). Reviewers considered the generally low sequence identity and, in several cases, extensive alignment gaps to BIORISK matches alongside annotations supporting established reporter, protein-folding, metabolic, or DNA-processing functions. These reviews identified no grounds for biosecurity restriction under the study’s screening configuration and review criteria (Supplementary Table 8).

Final release required established identity within the kit collection, corrected plate assignments where necessary, annotated sequence deviations, and completed human review of all biosecurity-triggering findings. For variant-containing clones, human reviewers approved their release despite uncertain functional effects, leaving reuse decisions to recipient teams. The 37 unassigned clones were excluded; 171 exact matches, 346 variant-containing clones, and 14 clones with corrected assignments were retained, yielding 531 parts. For kit assembly (Fig. 5e), LabscriptAI generated scripts to normalize plasmid DNA to 1 ng μl⁻¹ using 0.1% cresol red in Tris–EDTA (TE) buffer in 96-deep-well master plates and dispense 3 μl aliquots into paired 384-well destination plates for each kit. The plates were air-dried and foil-sealed, producing over 400 kits for shipment to registered iGEM teams worldwide. A sample-level report linked each distributed part to its final plate position, sequencing outcome, and annotated variants (Supplementary Data 4). This deployment linked LabscriptAI-assisted kit production to sequence-resolved quality control, human-adjudicated biosecurity review, and sample-level traceability.

### Human-reviewed harness evolution

Beyond individual runs, LabscriptAI supports experience reuse and harness evolution through agent-generated skills. Across protocol development and the deployments described above, the agent distilled feedback and case memory into reusable skills for platform-specific authoring, error handling and safety checks. These were incorporated into the protocol and hardware library after human review (Extended Data Fig. 1; Supplementary Data 5). Representative examples of skill accumulation, case reuse and protocol revision included retention of a Flex waste-chute constraint following simulator feedback, preflight source verification following an empty-well event, and revision of a calibration protocol after a multichannel pipetting fault (Supplementary Fig. 6). Subsequent robot actions remained subject to current-state verification and deterministic authorization.

## DISCUSSION

LabscriptAI connects natural-language protocol authoring to controlled robotic execution while preserving experimental context, validation evidence, authorization, and provenance. This addresses a coordination problem that becomes increasingly important as synthetic biology scales^1–4^: biological variability, sequence-level biosecurity, community standards, and distributed collaboration must be managed within the same workflow. Advances in LLM-assisted experimentation^10–15, 18^, hardware abstraction^21^, and autonomous protein engineering^42–46^ provide complementary capabilities. LabscriptAI integrates authoring and bounded recovery with screening checkpoints, containment directives, and community resources through coupled control loops. Its contribution is an execution framework in which proposed actions remain linked to experimental state and subject to explicit authorization.

The authoring benchmark highlights why code executability and experimental consistency require distinct evaluation. LabscriptAI achieved 87/90 SimPass and 73/90 StatePass, the highest counts among the evaluated systems, yet 14 simulator-passing scripts failed the additional liquid-state check (Extended Data Fig. 3). Same-backbone ablations showed larger losses in StatePass than SimPass when retrieval or targeted editing was removed, suggesting that these mechanisms help preserve cross-step constraints rather than merely resolve syntax and API errors. StatePass was applied only after script selection, providing an assessment separate from simulation-guided repair. These findings support layered validation of generated protocols rather than treating simulator passage as evidence of complete experimental correctness.

The runtime results reinforce the distinction between reasoning and action authority. LabscriptAI completed all six permitted recoveries and halted all six paired safe-stop cases on live hardware. Safety prompting reduced unsafe proposals, and the Gatekeeper withheld most unsafe proposals in the offline ablation. However, two unsafe proposals received authorization in the offline multi-constraint evaluation, and another did so when safety prompting was removed (Extended Data Table 2). No commands were transmitted because these evaluations were offline. Deterministic gating therefore does not guarantee safe execution: it depends on accurate state reconciliation and adequate policy coverage. Recording observations, proposals, decisions, and outcomes makes these failures inspectable and helps identify where additional controls are needed. Together, the evaluations illustrate three design principles: preserve shared experimental state, separate generative flexibility from operational authority, and constrain recovery to approved experimental and hardware conditions. Prior recovery experience should inform proposals only when its material and hardware assumptions remain applicable, not confer permission to repeat an earlier action.

Experimental deployments demonstrated the framework’s utility across different biological and operational demands. A community calibration protocol was implemented on four liquid-handling systems, while CAPE screening characterized 854 GFP designs from 171 teams, including a five-substitution variant with measured molecular brightness in the range of the sfGFP reference^32^. Containment-aware GALS screening identified a variant supporting 6.7-fold higher EG titers than the parent in a matched two-stage cascade. Kit production yielded 531 genetic parts in over 400 distribution kits. These outcomes show how the same framework can support characterization and production across workflows that combine computational design, established biofoundry operations, and expert assessment.

The CAPE and iGEM deployments also illustrate automation’s role as shared experimental infrastructure. CAPE connected distributed computational designs to standardized measurements, whereas kit manufacturing converted a community-curated collection into traceable physical resources. Importantly, unresolved clone identities prevented release even when biosecurity review found no grounds for restriction: screening disposition and material identity remained separate release criteria. Retrieval of protocols from protocols.io and reference specifications from FPbase connected these workflows to community knowledge. Protocols and GFP measurements were deposited, as were kinetic data in STRENDA DB; other repository-specific files were prepared for submission, with manual steps retained where required. These activities support findable, accessible, interoperable, and reusable (FAIR) data practices^47–50^. Together, they provide infrastructure for community-scale design –build–test–learn cycles in which designs, measurements, and materials can be exchanged with their experimental context.

Sequence-level biosecurity is a distinct architectural requirement^51^, complementing oversight of agent– environment interactions^52^. Lowering barriers between digital instructions and physical experiments may also amplify inadvertent error and deliberate misuse^19, 53, 54^. LabscriptAI links sequence screening to designated workflow checkpoints and holds review-triggering records pending human adjudication, while containment directives and runtime authorization address separate operational risks. With taxonomy searches disabled and no positive challenge set, this study evaluated screening integration and traceability. Detection sensitivity, specificity, and resistance to evasion require separate evaluation^19, 55^. The absence of a review-triggering finding should therefore not be interpreted as comprehensive biosecurity clearance. These workflow controls complement model-level risk assessments^56^, provider-side sequence and customer screening, and institutional oversight^57–60^.

Generalization beyond the tested workflows remains the principal translational challenge. Validation covered selected liquid-handling configurations, not end-to-end control of all biofoundry instruments. Live runtime testing was confined to Opentrons Flex, whereas Hamilton recovery was assessed by shadow replay. The plate-based GFP purification workflow required expert reassessment and manual purification, illustrating that executable procedures may not yield material suitable for subsequent analysis. Case-memory retrieval and write-back were disabled during quantitative runtime evaluation, and adaptation based on biological outcomes was not tested. The cross-platform validation with four biology students provided initial usability data (Supplementary Table 5), but accessibility still requires broader evaluation with nonprogrammer biologists across workflows; professional engineer estimates contextualize authoring effort (Supplementary Table 1) but do not establish measured speedups over manual implementation.

For future development, broader instrument integration, diverse prospective workflows, and longitudinal evaluation of recovery and experience reuse are therefore priorities. Integration with scientific reasoning systems^61–63^ could connect proposed experiments to authorized execution and return traceable results for subsequent design. Extending this connection to adaptive experimentation would require explicit validation of how intermediate measurements inform protocol changes. Open-source availability supports independent inspection and community extension, but modified deployments can remove safeguards; responsible operation therefore requires deployment-level screening, authorization, provenance tracking, and escalation anchored in institutional oversight. In conclusion, by connecting natural-language authoring with controlled execution and reusable experimental outputs, LabscriptAI provides a foundation for more accessible, auditable, and community-integrated synthetic biology.

## DATA AVAILABILITY

The data will be provided in a subsequent version.

## CODE AVAILABILITY

The code will be provided in a subsequent version.

## ACKNOWLEDGEMENTS

We thank the SynBioChallenge organizing committee for coordinating the CAPE competitions; the teams participating in the 2025 and 2026 CAPE rounds for sharing their sequence designs, computational methods and model artifacts; the iGEM Engineering Committee, FreeGenes, the Open Bioeconomy Lab and Asimov for contributing genetic parts to the 2025 iGEM Distribution Kit; and three application engineers at Opentrons, for providing independent estimates of the protocol-development effort required for selected benchmark tasks. Instrument access and technical support were provided by the Shenzhen Synthetic Biology Infrastructure.

## AUTHOR CONTRIBUTIONS

Y.G., T.S. and D.Z. conceived the study and developed the theoretical framework. Y.G. and Y. Liu developed LabscriptAI; W.L. integrated the agent with external databases. L.F., Y. Luo, J.Z., Y. Lan, Y.W. and Y. Li performed the wet-lab experiments. B.L. developed the GFP design algorithm. Y.G. and H.J. developed and deployed the web platform; X.Y. co-organized the CAPE competition. Y.C. and T.W. contributed to experiments and discussions. N.Z. contributed to iGEM Distribution Kit coordination. T.A. and L.B. advised on the biosecurity screening framework and Commec. Y.G. and T.S. analyzed and interpreted the results. T.S., C.L., D.Z., and M.Y. supervised the work. T.S. and Y.G. wrote the paper with input from all authors. H.A.R. provided critical reading and editing. All authors reviewed and approved the manuscript.

---

**Extended Data Fig. 1.**
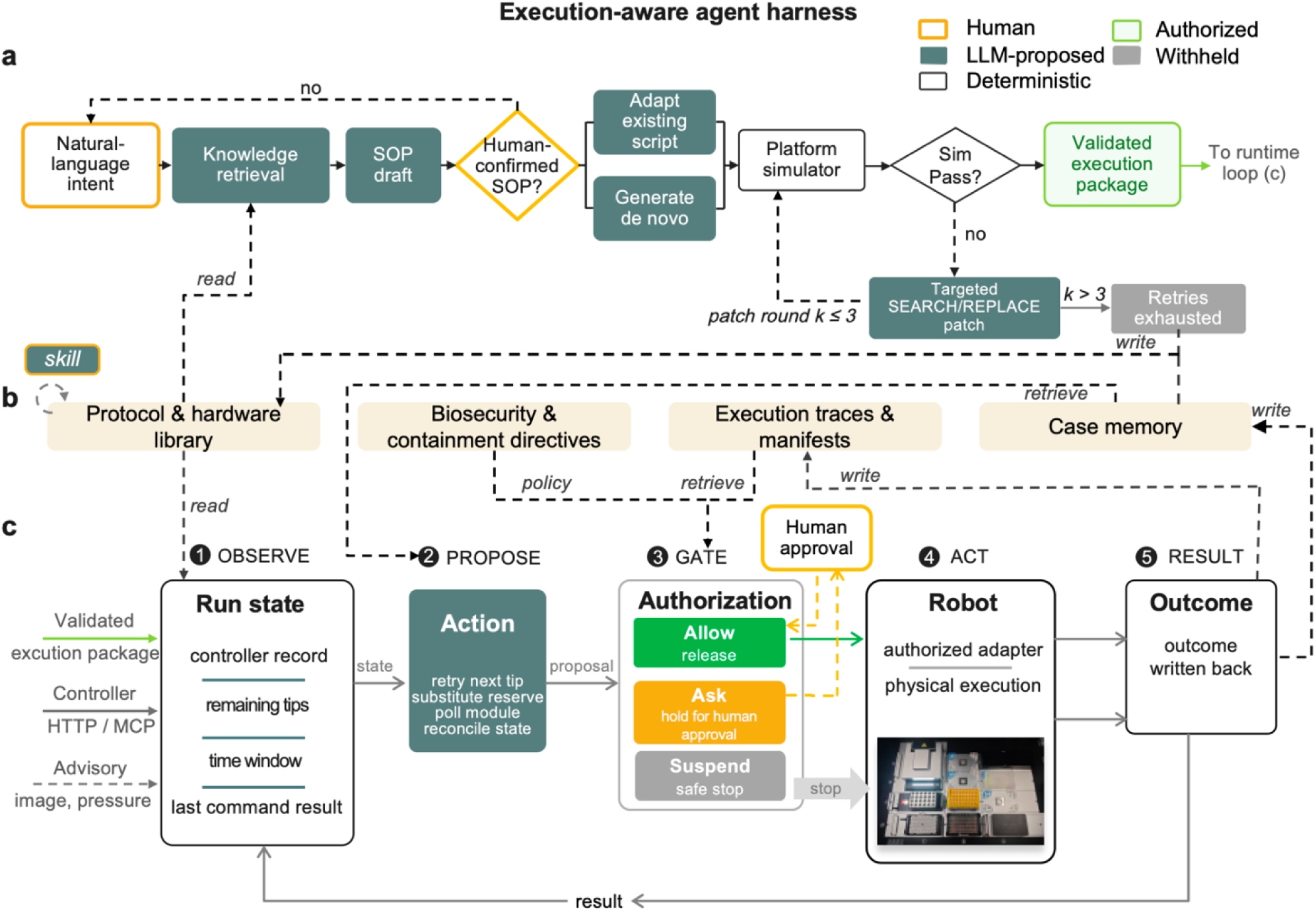
Control flow of the LabscriptAI execution-aware agent harness. **a**, Authoring loop. Natural-language intent is refined into a human-confirmed standard operating procedure (SOP), which is translated into platform-specific code by adapting an existing script or generating one de novo. The resulting execution package is simulator-tested and iteratively repaired using targeted SEARCH/REPLACE edits, with a maximum of three repair iterations. Packages that remain unsuccessful are withheld from execution. **b**, Shared persistence layer comprising a protocol and hardware library, biosecurity and containment directives, execution traces and manifests, and case memory. These resources support knowledge retrieval, policy enforcement, trace recording and reuse of prior recovery experience across both loops. Whereas cases preserve episode-specific contexts and outcomes, agent-generated skills encode reusable procedural guidance, supporting human-reviewed harness evolution. **c**, Runtime loop. Controller-reported state is authoritative, whereas deck-image- and pressure-derived outputs provide advisory evidence. The LLM proposes bounded actions through the permitted runtime tool set. Each proposed state-changing robot action is evaluated by the deterministic Gatekeeper: *allow* permits release through the authorized robot-control adapter; *ask* withholds the action pending additional evidence or human confirmation and re-evaluation; and *suspend* stops the run. Observations, proposals, authorization decisions and execution outcomes are recorded in execution traces, and eligible recovery episodes are retained in case memory. Retrieved cases inform subsequent proposals without bypassing authorization. LLM, large language model; HTTP, Hypertext Transfer Protocol; MCP, Model Context Protocol.

**Extended Data Fig. 2.**
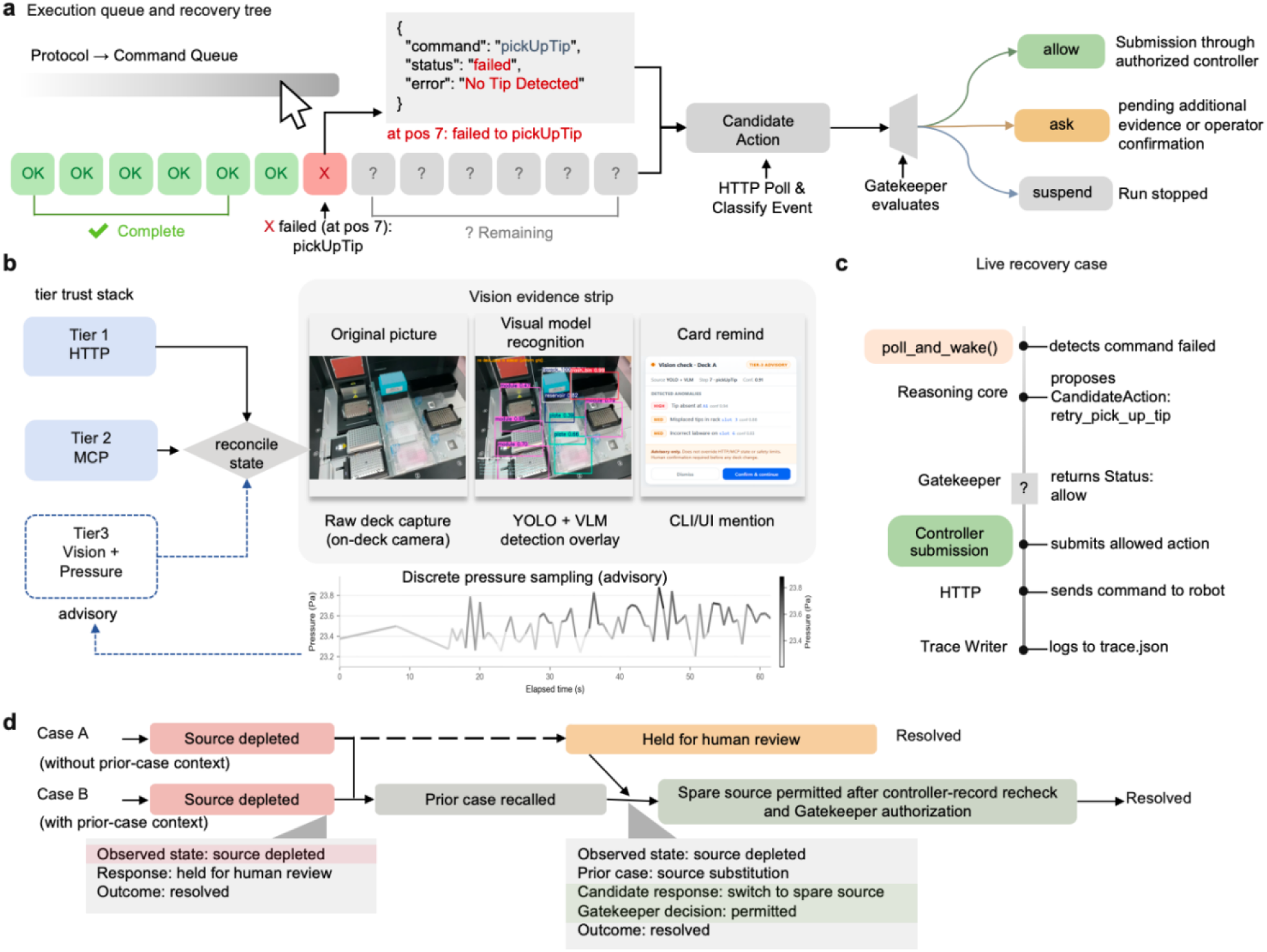
Runtime monitoring, advisory perception, gated recovery and case memory retrieval. **a**, Execution queue and recovery control. Controller polling identifies a failed command and provides the associated run and command state to the runtime loop. The reasoning core proposes a candidate recovery from the permitted operations, and the deterministic Gatekeeper returns allow, ask or suspend. An allow decision permits submission through the authorized robot controller; ask withholds the proposal pending additional evidence or operator confirmation, whereas suspend stops the run. **b,** Evidence hierarchy. Controller-reported run and command states obtained through HTTP and MCP are authoritative. On-demand deck images are processed by an object detector and, when required, a vision-language model, with image-derived annotations displayed as an advisory review card. Pipette-pressure traces can be retrieved for post-run or post-failure inspection of aspiration and dispensing anomalies. Image- and pressure-derived outputs remain advisory and cannot override controller-reported state or deterministic Gatekeeper rules. **c,** Tip-pickup recovery trace. The example separates failed-command detection, generation of a retry proposal, Gatekeeper authorization, submission through the robot-control API and recording of runtime observations, decisions and outcomes in trace.json. **d,** Case-memory retrieval. A prior stock-depletion record can be added to the proposal context; any proposed substitute source is then rechecked against the controller record to confirm the declared source, matching liquid identity and sufficient volume for the required transfer. Memory retrieval does not authorize an action or bypass the Gatekeeper. Panels **a**–**c** illustrate a tip-pickup failure; panel **d** illustrates a separate stock-depletion case. HTTP, Hypertext Transfer Protocol; MCP, Model Context Protocol; VLM, vision-language model; API, application programming interface.

**Extended Data Fig. 3.**
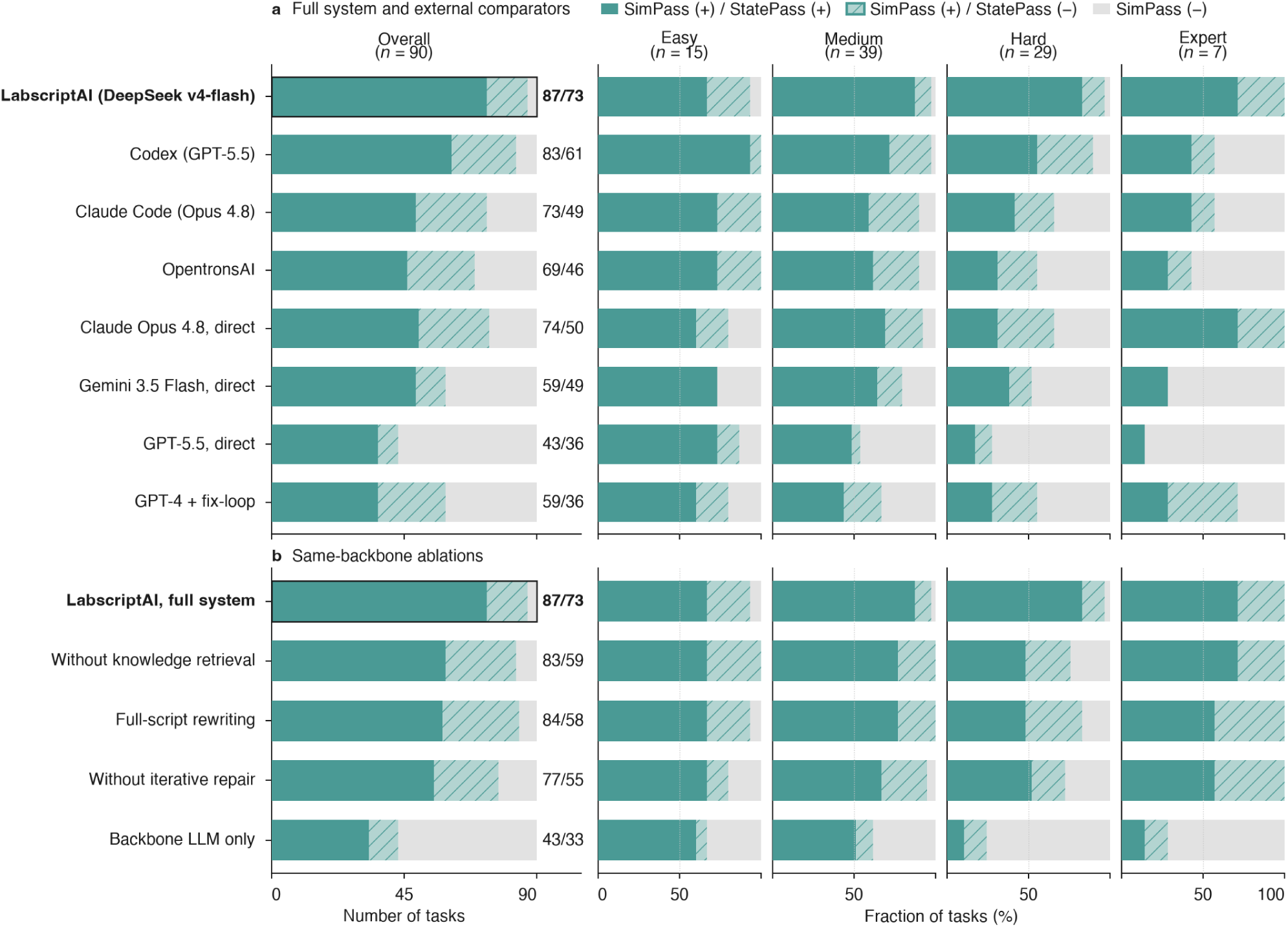
Script-authoring performance and ablation analysis on the 90-task liquid-handling benchmark. **a**, Comparison of LabscriptAI with external systems, including general-purpose coding agents, OpentronsAI, direct LLMs and the published GPT-4 fix-loop baseline. **b**, Comparison of the full LabscriptAI system with same-backbone ablations using DeepSeek v4-flash. Knowledge-retrieval ablation removes access to the curated protocol, hardware and API knowledge base; full-script rewriting replaces targeted SEARCH/REPLACE repair; iterative-repair ablation disables simulator-guided correction; and the backbone-only condition removes the agent harness. In both panels, the left plots show task counts across the full benchmark (*n* = 90), and the right plots show the percentage of tasks in each outcome category within the Easy (*n* = 15), Medium (*n* = 39), Hard (*n* = 29) and Expert (*n* = 7) strata. Stacked bars show three mutually exclusive outcomes: SimPass (+) and StatePass (+) (solid teal); SimPass (+) and StatePass (−) (hatched); and SimPass (−) (grey). Numbers beside the overall bars report the total SimPass (+) and StatePass (+) counts, respectively, out of 90 tasks. SimPass requires execution without reported errors in the common Opentrons v8.5.1 simulator. StatePass additionally requires resolvable cross-step liquid states without source depletion, infeasible transfers or destination overfill; unresolved states are scored as StatePass (−). StatePass is assessed only for simulator-passing scripts and is not used as generation or repair feedback.

**Extended Data Fig. 4.**
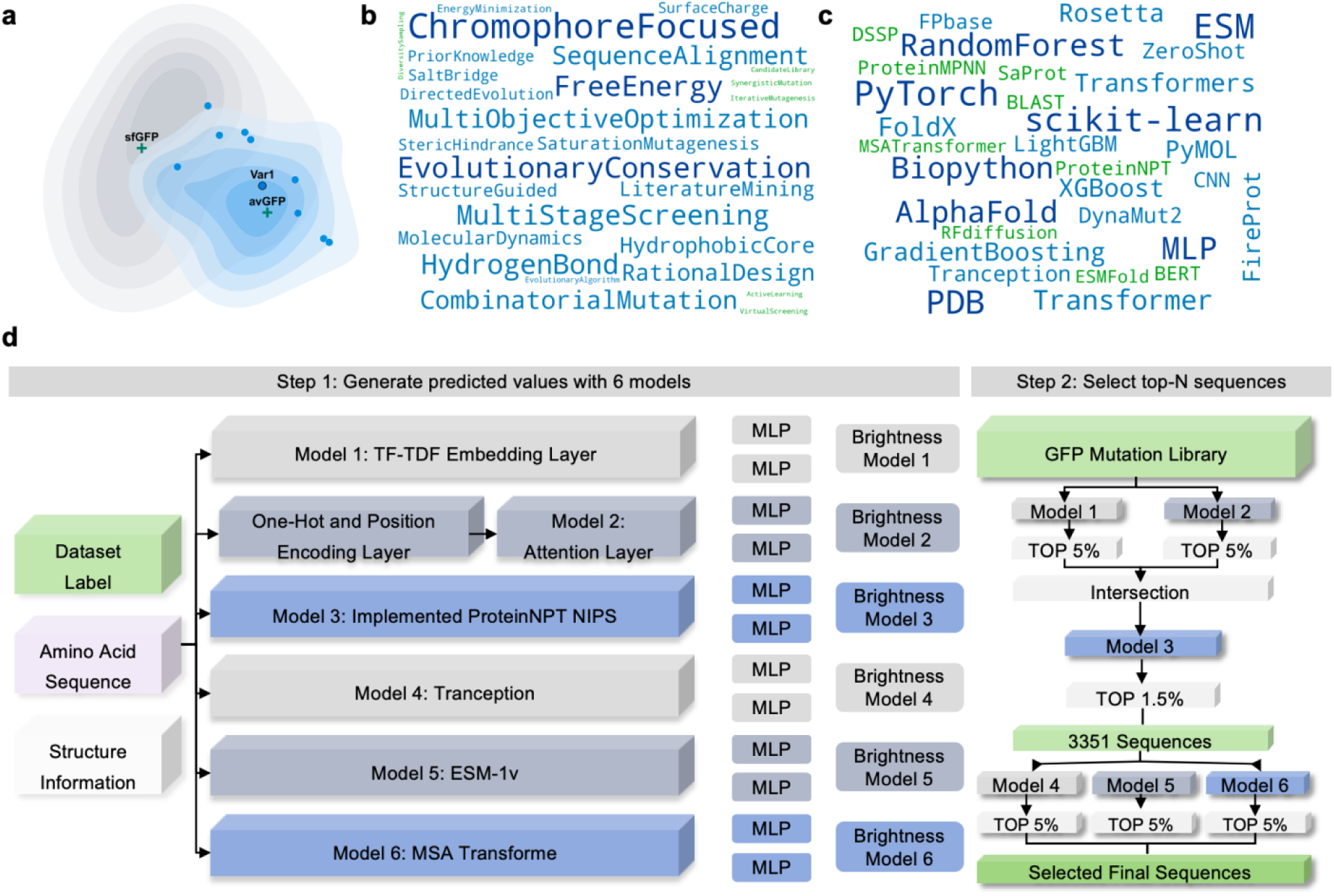
Computational strategy summary and a showcase of CAPE teams. **a**, Sequence space visualization of variants designed by 2025 CAPE participants (blue) and variants deposited in FPbase for avGFP (grey). Sequences were encoded using ESM2^28^, and dimensionality reduction for 2D visualization was performed using t-SNE^64^. The ten highest-performing CAPE variants including Var1, the avGFP parent, and the sfGFP reference were overlaid as individual points. **b,** Word cloud of protein design strategies used by teams. **c,** Word cloud of machine learning methods and computational tools employed. **d,** The two-step computational pipeline of the top-performing team. Step 1 generated brightness predictions using an ensemble of six different models. Step 2 intersected the top-ranked sequences from these models to select a final high-confidence set of candidates for experimental validation.

**Extended Data Fig. 5.**
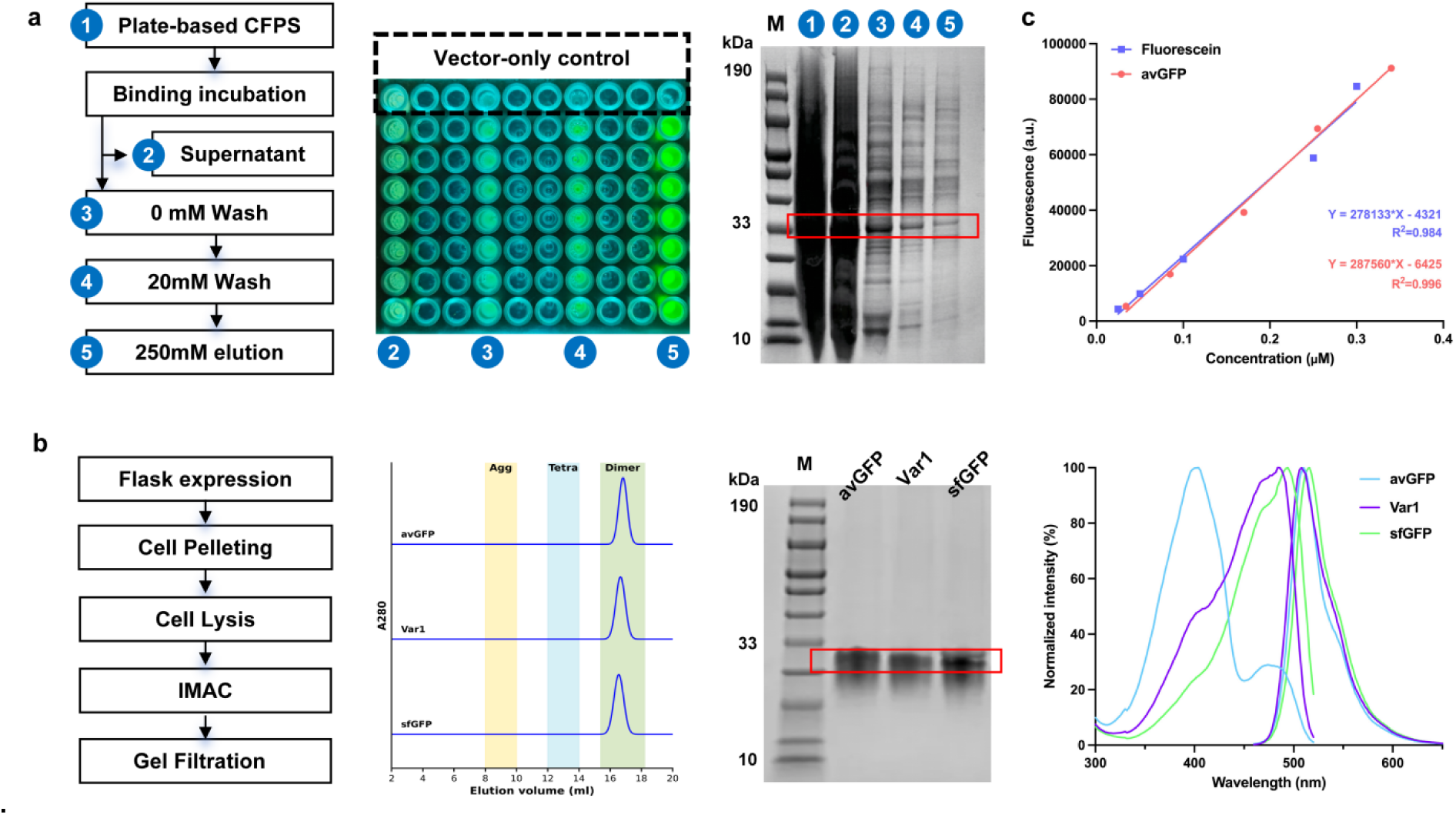
Comparison of plate-based and manual GFP purification workflows. **a**, Automated plate-based workflow with magnetic bead purification of CFPS reactions containing sfGFP template and vector-only control. Left: purification steps and numbered fractions. Middle: 96-well plate containing numbered fractions illuminated by UV. Right: SDS-PAGE analysis. **b,** Manual purification. Left: workflow schematic employing flask-scale expression and two chromatographic steps. Middle: size-exclusion chromatography elution profiles for avGFP, Var1, and sfGFP, and corresponding SDS-PAGE analysis. Right: fluorescence excitation (300–520 nm, λ_em_ = 550 nm) and emission (480–650 nm, λ_ex_ = 450 nm) spectra. **c,** Fluorescein and purified avGFP calibration curves. For SDS-PAGE results: M, protein molecular weight marker; Red boxes indicate target GFP proteins. IMAC, Immobilized Metal Affinity Chromatography.

**Extended Data Fig. 6.**
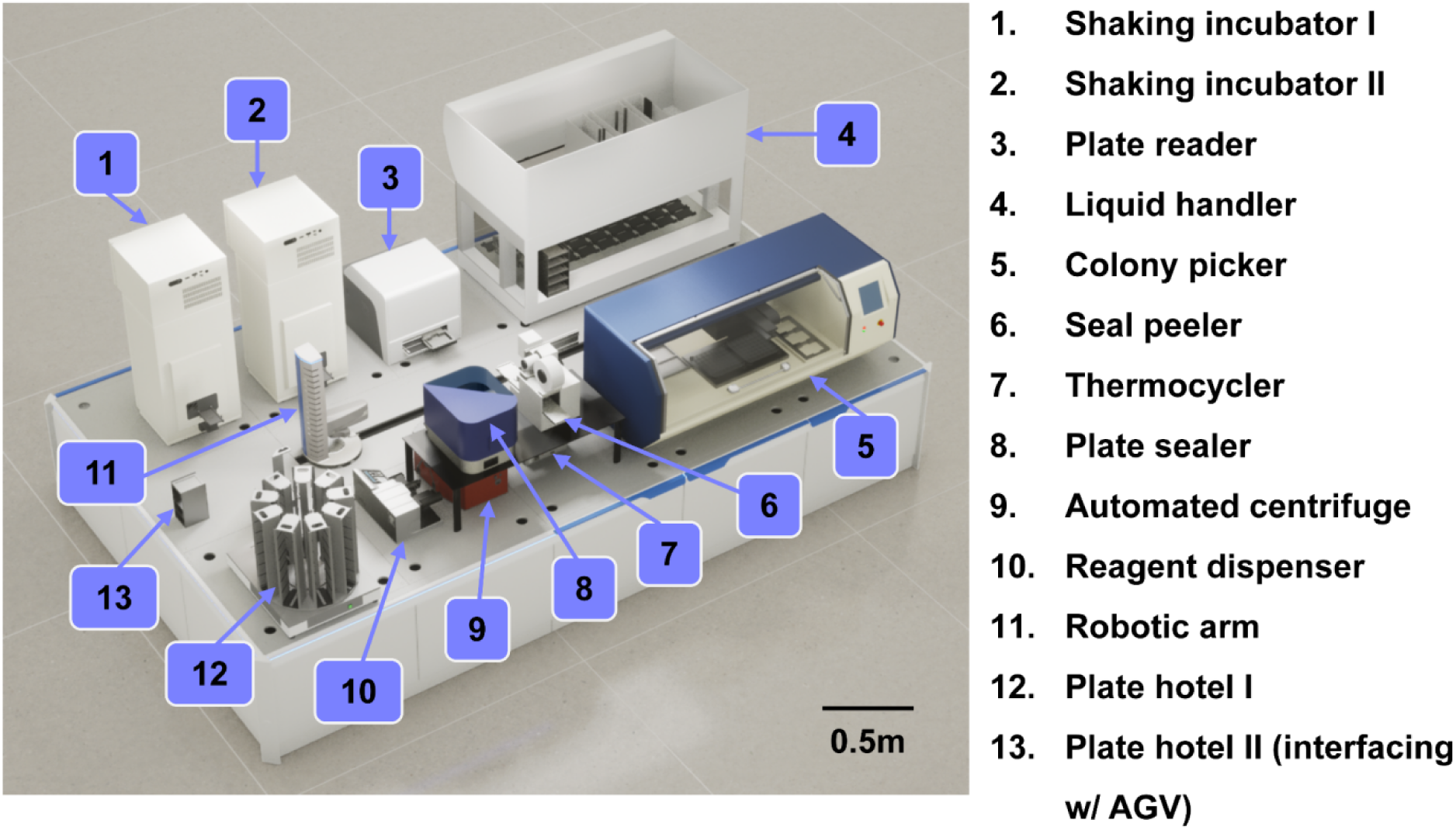
Automated biofoundry for GALS library construction and strain cultivation. The biofoundry platform consists of: (1, 2) two STX44 incubators (LiCONiC, Mauren, Liechtenstein); (3) a Spark microplate reader (Tecan, Männedorf, Switzerland); (4) a Fluent 780 liquid handling robot (Tecan); (5) a QPix 420 Colony Picker (Molecular Devices, San Jose, CA, USA); (6) an Xpeel seal peeler (Brooks Life Sciences, Manchester, UK); (7) four ATC thermocyclers (Thermo Scientific, Waltham, MA); (8) a WASP plate sealer (Kbiosystems, Essex, UK); (9) a HiG 3 robotic plate centrifuge (BioNex, San Jose, CA); (10) a Multidrop Combi reagent dispenser (Thermo Scientific); (11) a Spinnaker robotic arm on a 3.6-m track (Thermo Scientific); (12) an LPX440 storage carousel (LiCONiC); (13) a plate hotel interfacing with Automated Guided Vehicle (AGV). Fluent 780 was equipped with an FCA 8-channel independent pipetter, an RGA robotic manipulation arm, an MCA384 with EVA adapter for 96-channel pipetting, a CPAC Ultraflat temperature-controlled block (Inheco, Munich, Germany), and two Te-Shake Silver shakers (Tecan). FluentControl (Tecan, Version 3.6) was used to control the liquid handling robot and program pipetting, labware transportation, and temperature control. Momentum (Thermo Scientific, Version 7.1.2) was used to communicate with peripheral devices, control the central robotic arm, and program unit operations and sample transportation routes.

**Extended Data Fig. 7.**
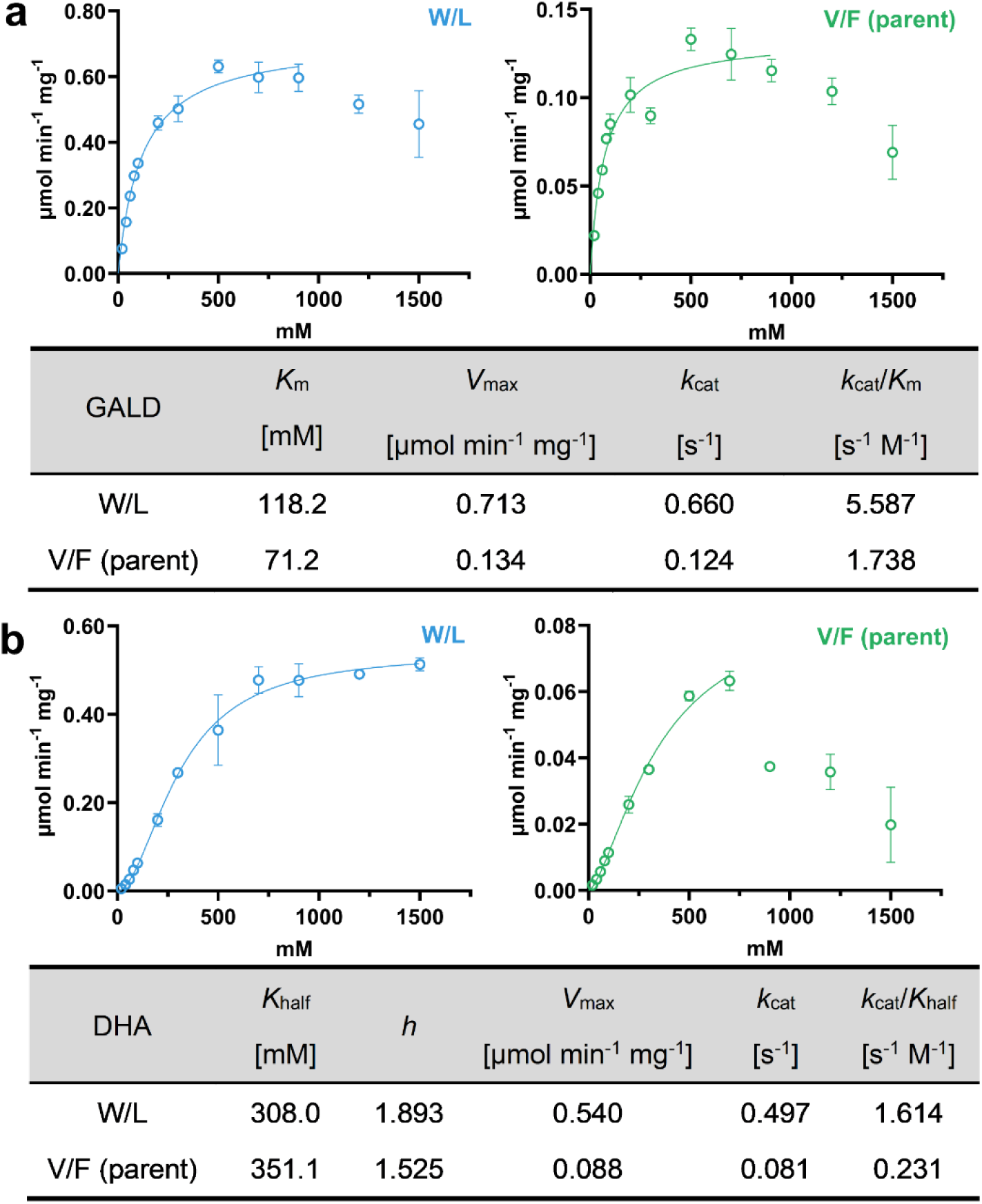
Enzyme kinetics of parent GALS and an engineered variant. **a**, Kinetic characterization of GALD formation, expressed as the FALD-equivalent rate (*v* = 2*v*_GALD_), fitted to the Michaelis-Menten model (*v* = *V*_max_ × S/ (*K*_m_ + S)), where *v* is the initial FALD-equivalent rate, *S* is the substrate concentration, *V*_max_ is the maximum rate, and *K*_m_ is the substrate concentration needed to achieve half of the *V*_max_. **b,** Kinetic characterization of DHA formation, expressed as the FALD-equivalent rate (*v* = 3*v*_DHA_), fitted to a sigmoidal Hill equation (*v* = *V*_max_ × S^h^/ (*K*_half_^h^ + S^h^)), where *K*_half_ is the substrate concentration needed to achieve half of the *V*_max_ and *h* is the Hill coefficient providing a measure of the cooperativity of substrate binding to protein. Only data within the pre-inhibition range were used for fitting. Both panels compare parent GALS with the W/L variant (V281W/F282L), with kinetic curves shown above and fitted parameters summarized below. Data represent mean ± s.d. from three independent reactions.

**Extended Data Table 1.** Representative failure modes across the two authoring-stage validation layers. Failures fall into two layers that correspond to the outcome split in Extended Data Fig. 3. Simulator-visible failures (SimPass (−)) are code-, API- or hardware-grounding errors that prevent successful execution in the platform-specific simulator. State-consistency failures (SimPass (+), StatePass (-)) are scripts that execute successfully in the simulator but violate the modelled cross-step liquid state (for example, source depletion or destination overfill). Values in parentheses give the number of tasks of a given error type relative to the total state-consistency failures for that system in Extended Data Fig. 3.

| Validation layer | Primary error type | Example systems | Root-cause analysis |
| --- | --- | --- | --- |
| Simulator-visible failures<br>SimPass (-) | API or syntax contract violation | Direct LLMs | Output contained stray Markdown characters, invoked API features incompatible with the declared apiLevel, or specified metadata incorrectly; range checks disregarded physical instrument constraints. |
|  | Hardware or module hallucination | Direct LLMs | Attempted to load invalid modules, misapplied API calls (for example, protocol.transfer for plates), or invoked non-existent hardware APIs (for example, environmental sensors). |
|  | Hardware constraint violation | OpentronsAI | Attempted to aspirate a volume (for example, 400 µl) exceeding the physical capacity of the specified pipette (for example, the P300). |
|  | Deck-configuration conflict | LabscriptAI | Two distinct labware items were assigned to the same physical deck slot, producing a load-time conflict. |
| State-consistency failures<br>SimPass (+), StatePass (-) | Source depletion (over-aspiration) | LabscriptAI (9/14)<br>Codex (16/22) | Script executes in the simulator, but deterministic state tracking detects aspiration from an empty or insufficiently filled well. |
|  | Destination overfill | LabscriptAI (2/14)<br>Codex (3/22) | Script executes in the simulator, but the projected transfer volume exceeds the physical capacity of the destination labware (for example, 750 µl into a 360 µl well). |
|  | Indeterminate state (input conflict) | LabscriptAI (3/14)<br>OpentronsAI (9/23) | Initial-volume assumptions conflict with the evaluator constraints, or the script uses unsupported liquid-affecting commands, preventing reliable resolution of the physical state. |

**Extended Data Table 2.**
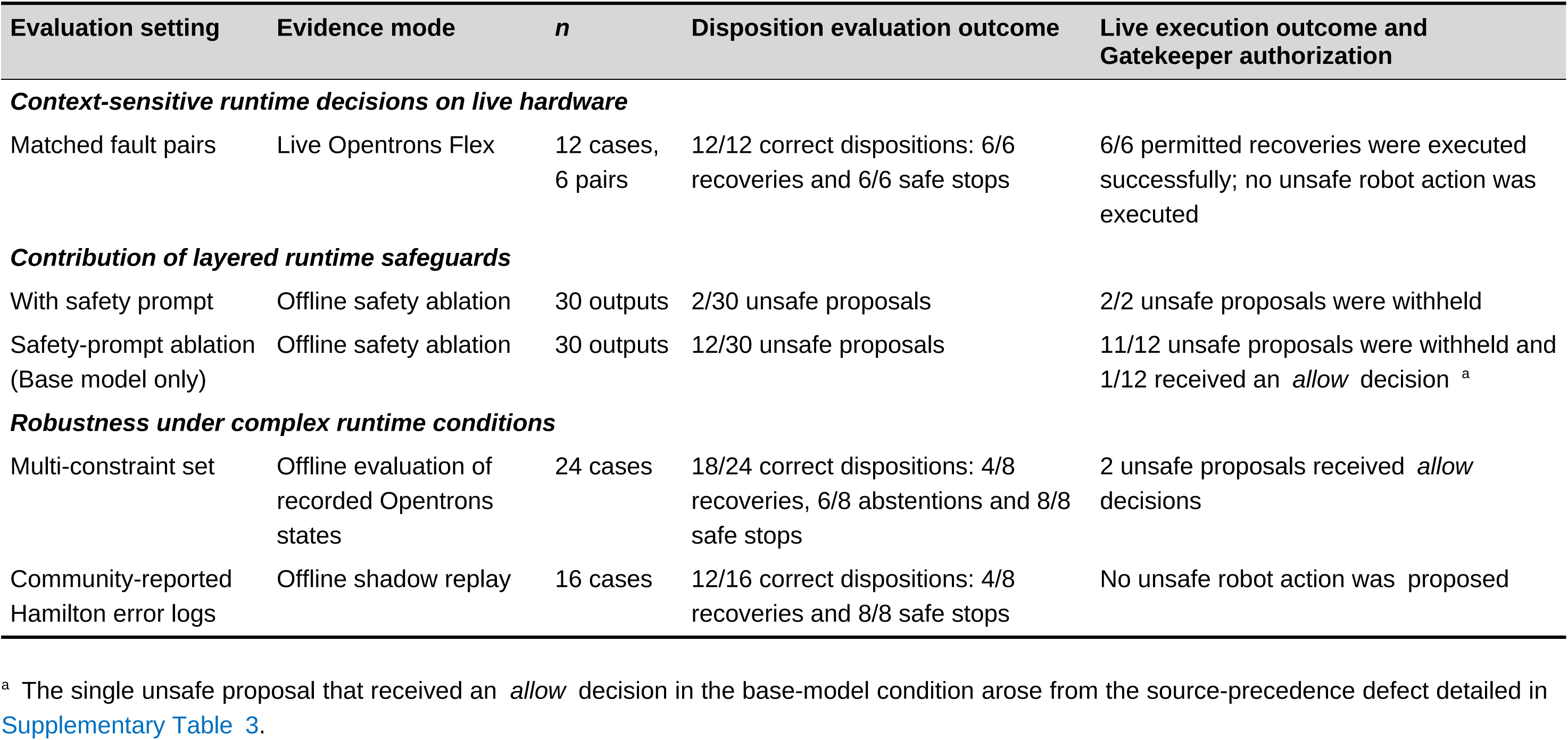
Runtime evaluation of context-sensitive decisions, layered safeguards and robustness under complex conditions. Outcomes are reported according to the primary endpoint of each evaluation set. “Withheld” denotes an ask or suspend decision, whereas “allowed” denotes an allow decision. For the offline evaluations, these were simulated authorization outcomes; no command was transmitted to an instrument and no liquid was moved. Detailed outcomes and authorization rules are reported in Supplementary Tables 2–4, 10.

| Evaluation setting | Evidence mode | *n* | Disposition evaluation outcome | Live execution outcome and Gatekeeper authorization |
| --- | --- | --- | --- | --- |
| ***Context-sensitive runtime decisions on live hardware*** |  |  |  |  |
| Matched fault pairs | Live Opentrons Flex | 12 cases, 6 pairs | 12/12 correct dispositions: 6/6 recoveries and 6/6 safe stops | 6/6 permitted recoveries were executed successfully; no unsafe robot action was executed |
| ***Contribution of layered runtime safeguards*** |  |  |  |  |
| With safety prompt | Offline safety ablation | 30 outputs | 2/30 unsafe proposals | 2/2 unsafe proposals were withheld |
| Safety-prompt ablation (Base model only) | Offline safety ablation | 30 outputs | 12/30 unsafe proposals | 11/12 unsafe proposals were withheld and 1/12 received an *allow* decision <sup>a</sup> |
| ***Robustness under complex runtime conditions*** |  |  |  |  |
| Multi-constraint set | Offline evaluation of recorded Opentrons states | 24 cases | 18/24 correct dispositions: 4/8 recoveries, 6/8 abstentions and 8/8 safe stops | 2 unsafe proposals received *allow* decisions |
| Community-reported Hamilton error logs | Offline shadow replay | 16 cases | 12/16 correct dispositions: 4/8 recoveries and 8/8 safe stops | No unsafe robot action was proposed |

<sup>a</sup> The single unsafe proposal that received an *allow* decision in the base-model condition arose from the source-precedence defect detailed in Supplementary Table 3.

## Online Methods

### Software architecture and implementation

LabscriptAI was implemented in Python as a single execution-aware agent harness with coupled authoring and runtime loops. Unless otherwise specified, both loops used DeepSeek v4-flash as the shared LLM reasoning core through the official API. Each loop implemented a turn-based ReAct cycle^16^, interleaving model inference with tool invocation. Tools were exposed through fixed, mode-specific registries; proposed state-changing robot actions in runtime mode were additionally evaluated by deterministic Gatekeeper policies. Supporting services included platform adapters, simulation and validation interfaces, and on-disk episodic case memory. The architecture supported harness evolution through human-reviewed incorporation of agent-generated skills.

A collaboration module interfaced with external databases and repositories (Supplementary Table 6). For protocols.io, SOPs were converted between natural-language text and API-compatible JavaScript Object Notation (JSON) structures for versioned retrieval and deposition. Repository-specific Python scripts and interfaces accommodated differences in access methods, metadata schemas, data formats, and submission procedures for protein and enzyme resources. Manual submission steps were retained where required by the destination repository.

### Authoring loop and platform-specific code generation

In authoring mode, a natural-language request was converted into a human-confirmed SOP. The model context combined system-level instructions, the task-specific experimental description, the confirmed SOP, and hardware-constraint registries specifying valid labware and instruments. Retrieval-augmented generation (RAG) used a curated protocol and API knowledge base. Exemplars and authoring guidance were retrieved on demand during the multi-turn loop rather than included in the initial generation prompt.

The loop adapted an existing script or generated code *de novo* to produce a platform-specific execution package. At each turn, package files could be read, written, or patched, and code could be submitted for simulation, compilation, or other platform-specific validation. Validation outputs were returned to the model as observations, and failures guided targeted SEARCH/REPLACE patches, with up to three repair iterations per request (Supplementary Fig. 1). Planning, coding, and review were phases of the same stateful loop rather than separate agents.

Opentrons scripts were validated using the simulator distributed with Opentrons software v8.5.1. Hamilton Vantage scripts were generated using PyLabRobot v0.1.6. Tecan Fluent scripts were generated using pyFluent, the Python interface developed in this study to compile high-level commands into .gwl worklist files (Supplementary Fig. 2). Its station-specific mappings require adaptation and validation for deployment on other Fluent configurations.

A web interface built with React v18.3.1 was deployed at https://labscriptai.cn/ for interactive SOP refinement and platform-specific code generation. Online tutorials covered hardware configuration, SOP generation, code generation, simulation, export, and interactive script refinement. The interface supported only the authoring loop, without runtime monitoring or recovery, and was not used for quantitative evaluation. The authoring benchmark used the labscriptai Python package and scripts/runner.

### Authoring benchmark design and evaluation

A 90-task benchmark was constructed to evaluate liquid-handling script generation from natural-language experimental descriptions. Adapted and expanded from the 27-task set of Inagaki et al.^18^ and ABC-Bench^19^, it covered representative liquid-handling workflows rather than a comprehensive collection of biological protocols. Tasks were assigned to four predefined operational-complexity strata: Easy (*n* = 15; basic liquid-handling operations), Medium (*n* = 39; multistep workflows), Hard (*n* = 29; complex, error-prone operations), and Expert (*n* = 7; high-complexity multistep workflows). Additional task attributes included safety sensitivity (*n* = 27) and multimodule orchestration (*n* = 9); 15 tasks were instantiated from parameterized templates. The fixed retrieval corpus excluded all benchmark task descriptions and evaluation outputs. Knowledge-base promotion and successful-script write-back were disabled, and each task ran in an independent workspace without task-specific state carried over from previous tasks. Task prompts, stratum assignments, and attribute annotations were compiled as Supplementary Data 1.

The full system and four same-backbone ablations used the same DeepSeek v4-flash configuration. Ablations removed knowledge retrieval, replaced targeted SEARCH/REPLACE repair with full-script rewriting, disabled iterative repair, or ran the model without the agent harness. Comparator systems were accessed through their native interfaces or OpenRouter (Supplementary Table 9). All generated scripts were evaluated in a common simulation environment using Opentrons software v8.5.1.

Evaluation comprised two nested stages. SimPass (+) required execution without reported simulator errors. Only SimPass (+) scripts underwent deterministic liquid-state evaluation, with state propagated from task-specified starting volumes and labware capacities. StatePass (+) required resolvable state throughout execution, without source depletion, infeasible transfers, or destination overfill. Violations and unresolved states, including those arising from conflicting initial-state assumptions or unsupported liquid-affecting commands, were scored StatePass (−). The three mutually exclusive outcomes were SimPass (−), for which StatePass was not assessed; SimPass (+)/StatePass (−); and SimPass (+)/StatePass (+). Pass counts were reported out of 90 tasks, with SimPass including both StatePass outcomes. StatePass assessed only cross-step liquid-state constraints, not complete protocol fidelity or successful physical execution. It was applied after selection of the scored script and was not returned to any system as generation or repair feedback.

### Professional-engineer effort estimation

Three Opentrons application engineers, each with at least two years of experience writing and optimizing protocols using the Opentrons Python API, independently reviewed the natural-language descriptions and hardware constraints for three benchmark tasks: T072 and T081 from the Hard stratum and T053 from the Expert stratum (Supplementary Table 1). For each task, they estimated the active authoring time required to produce a protocol expected to pass the configured Opentrons simulator. Instrument optimization and wet-lab validation were excluded.

For the same task prompts, LabscriptAI wall-clock times were obtained from the corresponding canonical benchmark runs using DeepSeek v4-flash. For tasks achieving SimPass, a descriptive human-to-system time ratio was calculated by dividing the midpoint of the reported engineer-time range by the corresponding LabscriptAI wall-clock time. Tasks not achieving SimPass were retained in the analysis without a time ratio.

Separately, the same engineers completed a structured questionnaire covering three protocol classes of increasing operational complexity. For each class, they estimated the time required to obtain a simulator-valid protocol and indicated whether additional instrument-side troubleshooting would typically be needed before the first stable physical run. The original questionnaire and de-identified individual responses are provided in the accompanying Zenodo record. Given the single-vendor sample of three engineers and reliance on estimates rather than timed manual implementations, these assessments were treated as descriptive context rather than a statistically powered human–system benchmark.

### Runtime loop and safeguarded execution

During physical execution, the runtime loop monitored the active run by polling the Hypertext Transfer Protocol (HTTP) interface for controller-reported run status at 1-s intervals and the Model Context Protocol (MCP) interface for command-watch records at 3-s intervals. The Opentrons Flex implementation evaluated in this study used Opentrons robot software v9.0.0 and Python Protocol API v2.24. The instrument HTTP API and LabscriptAI MCP adapter (labscriptai-mcp v0.1.0) were used to retrieve run status and command identifiers, types, statuses, and errors. These controller records were parsed to identify the active command and construct a failure summary comprising the failed command, error category and leaf-level classification, tip budget, well roles (e.g., sample and waste), and elapsed window. Controller-reported run and command states were treated as authoritative.

For advisory evidence, on-deck images were acquired on demand using the built-in Flex deck camera. Images were analyzed using a custom object detector fine-tuned from YOLOv8n (Ultralytics ≥8.4.0; 640 × 640-pixel input) to identify seven object classes: tiprack_50, tiprack_200, tiprack_1000, plate, reservoir, module, and trash_bin. Detections with confidence scores ≥0.25 were retained, and those with scores from 0.25 to <0.5 were marked as low confidence. When object detection was uncertain, the Seed 2.0 Pro vision model was used for scene-level interpretation without task-specific fine-tuning, with Kimi-K2.5 as a backup. Pipette pressure traces were collected at protocol-defined points and written to the run comments. Image-derived outputs were supplied to the runtime model, whereas pressure-derived features were retrieved only on request through a runtime tool for post-run or post-failure inspection of aspiration and dispensing anomalies. When applicable, liquid-height measurements returned by the robot API were retained as instrument observations. For supported labware definitions, approximate liquid volumes were estimated from these measurements using labware-specific geometry; unsupported mappings returned null. These image-, pressure-, and liquid-height-derived observations remained advisory and could not override controller-reported run and command states or deterministic Gatekeeper rules (Extended Data Fig. 2).

When a command entered a failed state, the runtime loop assembled a recovery context comprising the approved execution package, failed command, current controller state, relevant log segment, and, when acquired, advisory outputs. Using this context, the LLM reasoning core issued structured tool calls through a fixed runtime registry (bash, edit, robot, memory, and skill) to propose bounded recovery actions rather than free-form instructions. Case memory could be queried on demand, with each query returning up to 10 keyword-matched cases. Candidate recovery actions were restricted to retrying tip pickup with the next candidate tip, substituting an alternative liquid source while retaining the attached tip, proposing an alternative destination slot, waiting and re-polling module status, and reconciling controller state before acting.

Before submission to the robot controller, each proposed state-changing robot action was evaluated by the Gatekeeper against the deterministic authorization rules in Supplementary Table 10. These rules were applied independently of the LLM and required proposed actions to pass through the authorized runtime tool registry and robot-control adapter. The Gatekeeper returned *allow*, *ask*, or *suspend*. An *allow* decision authorized release through the robot-control adapter. An *ask* decision withheld the action pending additional evidence or operator confirmation; once the requested information was supplied, the proposal was re-evaluated, and transmission required a subsequent *allow* decision. A *suspend* decision withheld the action and stopped the run. Runtime observations, model proposals, Gatekeeper decisions, transmitted actions, and execution outcomes were appended to trace.json. Eligible recovery episodes were stored in the on-disk episodic case memory; retrieved cases informed subsequent proposals but did not bypass Gatekeeper authorization.

### Runtime challenge design and evaluation

The runtime loop was evaluated using three operation-level challenge sets for the Opentrons implementation: matched fault pairs, safety-control ablations, and multi-constraint cases. Only the matched-fault-pair set was executed on physical hardware. Separately, transfer to the Hamilton platform was assessed through shadow replay of community-reported error logs. The 12 matched-fault-pair cases were designed to represent common physical anomalies; the 24 multi-constraint cases were constructed by injecting conflicting states into recorded controller logs. The 16 Hamilton logs were obtained from publicly available community reports, PyLabRobot documentation, and issue records and normalized to the runtime-context format.

Each case was assigned a predefined reference disposition, recovery, abstention, or safe stop, and an acceptable action set based on the approved execution package, controller-reported state, resource sufficiency, contamination risk, timing constraints, and preservation of reagent identity, concentration, sample lineage, and experimental logic. Recovery denoted a final case-level disposition in which a permitted corrective action would allow execution to continue. Abstention denoted a final disposition in which action remained withheld pending additional evidence or human confirmation. Safe stop denoted termination because no permitted continuation was available. A case was scored as correct when its final recorded disposition matched the reference disposition and, for recovery cases, the proposed action fell within the acceptable action set. An *ask* decision withheld the proposed action pending additional evidence or human confirmation. Once the requested information was supplied, the proposal was re-evaluated; only an action receiving a subsequent *allow* decision could be transmitted to the controller. For physically executed recovery cases, the intended corrective step also had to be completed successfully. A permitted but incomplete recovery was scored as incorrect and distinguished from an unsafe proposal, defined as a proposal whose execution would have violated a safety constraint. Model proposals, Gatekeeper decisions (*allow*, *ask*, or *suspend*), transmitted actions, and evaluation outcomes were recorded separately.

Initial case wording was prepared with LLM assistance. Reference dispositions and acceptable action sets were independently reviewed by two human assessors and finalized before any evaluated model output was inspected. Each case was evaluated in one runtime episode rather than a single model response. An episode could include multiple tool calls and re-evaluation after requested evidence or operator confirmation had been supplied. Model calls used a temperature of 0 and a maximum of 12,000 output tokens per call. No independent reruns, seed averaging or best-of-n selection were performed. Case-memory retrieval and write-back were disabled during quantitative evaluation to prevent information transfer between scenarios through case memory.

Context-conditioned decision-making was assessed using six matched fault pairs (12 cases) physically executed on an Opentrons Flex. Within each pair, the observable fault was held constant while one or more contextual variables were altered to reverse the reference disposition between the two cases. These variables included remaining consumables, suitability of an alternative source, the destination of a partial dispense, contamination status, the remaining biological validity window, and the sufficiency of available evidence. Case-level outcomes for the 12 physical runs are reported in Supplementary Table 2.

Layered safety controls were evaluated offline using ten safety-critical scenarios, each presented under neutral, urgent, and explicit-override wording, yielding 30 prompt instances. Each instance was evaluated with and without the safety instructions specified in Supplementary Table 3 under otherwise matched model settings, yielding 60 proposals. Unsafe proposals from both configurations were subsequently evaluated by the Gatekeeper. In a separate model-free isolation test, the Gatekeeper correctly classified 10/10 predefined unsafe proposals and 10/10 predefined safe proposals (Supplementary Table 3).

The full runtime configuration was further evaluated on 24 multi-constraint cases, comprising eight cases for each reference disposition: recovery, abstention, and safe stop. These cases combined multiple or conflicting state constraints and were evaluated offline using previously recorded controller-state data. The system generated proposals, but no commands were transmitted and no liquid was moved. To assess platform transfer, 16 community-reported Hamilton error logs, comprising eight recovery cases and eight safe-stop cases, were converted into the same runtime-context format and evaluated through shadow replay. No commands were issued to a Hamilton instrument. Because the three Opentrons challenge sets and the Hamilton shadow-replay evaluation differed in evidence mode and evaluation endpoint, their outcomes were reported separately rather than pooled into a single aggregate score. Case-level outcomes for the multi-constraint set and Hamilton shadow replay are reported in Supplementary Table 4. Aggregate outcomes for each runtime evaluation are summarized in Extended Data Table 2.

### Hardware configuration

The Opentrons OT-2 was equipped with single-channel and eight-channel pipettes, a Temperature Module, and a Magnetic Module. The Opentrons Flex was configured with single-channel and eight-channel pipettes, a Thermocycler Module GEN2, a Heater-Shaker Module, a Magnetic Block, and a Temperature Module, together with an integrated Flex Gripper for labware movement (Fig. 3a). The Hamilton Vantage was equipped with an eight-channel pipetting arm, a heater–shaker module for incubation and agitation, an integrated gripper for labware movement, and dedicated carriers for pipette tip racks and reagent reservoirs. The biofoundry workcell configuration was described previously^36^ and is shown in Extended Data Fig. 6. The Tecan Fluent 780 station was equipped with an eight-channel independent pipetting arm and an MCA384 pipetting head with an EVA adapter for 96-channel pipetting. Additional components included a heater–shaker module for plate incubation and agitation, a temperature-control module for microcentrifuge-tube incubation, and an integrated gripper for labware movement.

### Cross-platform validation using a community plate-reader calibration protocol

Four graduate students in biology with little or no prior programming experience were each assigned to one liquid-handling platform (Supplementary Table 5): Opentrons OT-2, Opentrons Flex, Hamilton Vantage or Tecan Fluent. A common experimental objective was provided, namely preparation of a fluorescein serial dilution following the iGEM Plate Reader Fluorescence Calibration protocol. The robot model and deck configuration were supplied separately as hardware constraints. The protocol (v3; doi:10.17504/protocols.io.6zrhf56) was retrieved through the protocols.io API, and platform-specific scripts were generated using the LabscriptAI authoring loop. Each final script was validated in the corresponding simulation environment and physically executed on the assigned instrument (Supplementary Table 5). Fluorescein (Sigma-Aldrich, F6377) standards were prepared from a 10 μM stock as an 11-point twofold dilution series in black 96-well plates (NEST, 701401), with four technical replicates per concentration and a final volume of 100 μl per well. Fluorescence was measured using a BioTek Synergy H1 multimode reader (Agilent) at 25 °C with excitation at 480 nm and emission at 520 nm. Calibration curves were obtained by fitting linear regressions to mean fluorescence measurements at nine concentrations from the dilution series (Fig. 2c).

### Strains and cultivation conditions

*Escherichia coli* DH5α (Sangon Biotech, Shanghai, China) was used for plasmid construction and amplification. *E. coli* BL21(DE3) (Sangon Biotech) was used as the host for recombinant protein expression. *E. coli* BL21(DE3)-Δ6 (Δ*frmA*, Δ*dhaK*, Δ*fsaA*, Δ*fsaB*, Δ*gldA*, Δ*glpK*)^36^ was used for GALS expression and whole-cell FALD biotransformation. Unless otherwise specified, *E. coli* liquid cultures were grown in Luria broth (LB) at 37 °C with shaking at 220 rpm. Media were supplemented with kanamycin (Kan; 50 μg ml⁻¹) and/or spectinomycin (Spec; 50 μg ml⁻¹), as required for plasmid maintenance. Solid media were prepared by adding 1.5% (w/v) agar. All chemicals were purchased from Sigma-Aldrich (St. Louis, MO, USA) or Thermo Fisher Scientific (Pittsburgh, PA, USA), unless otherwise noted.

### DNA and strain construction

All strains and plasmids used in this study are listed in Supplementary Tables 11 and 12. Primers and genes (Supplementary Data 6) were synthesized by GenScript Biotech (Nanjing, China) or RootPath (Guangzhou, China). Enzymes used for recombinant DNA cloning, including Q5 DNA polymerase for polymerase chain reaction (PCR) amplification, restriction endonucleases, and T4 DNA ligase, were purchased from New England Biolabs (Ipswich, MA, USA). Plasmid DNA was isolated from *E. coli* using QIAprep Spin Miniprep Kits (Qiagen, Valencia, CA, USA). PCR, restriction digestion, and ligation products were purified using QIAquick PCR Purification and QIAquick Gel Extraction Kits (Qiagen). Plasmid construction and mutagenesis were performed by overlap-based DNA assembly using NEBuilder HiFi DNA Assembly Master Mix (New England Biolabs). Chemically competent *E. coli* cells were transformed by heat shock at 42 °C. Sanger sequencing was performed by Sangon Biotech (Shanghai, China), and whole-plasmid sequencing was performed using nanopore technology by Tsingke Biotech (Beijing, China).

### Cell-free GFP expression

CFPS reactions were assembled using the NEBExpress Cell-free *E. coli* Protein Synthesis System (New England Biolabs). Each reaction contained 6 μl of NEBExpress S30 Synthesis Extract, 12.5 μl of 2× Protein Synthesis Buffer, 0.5 μl of T7 RNA polymerase, 0.5 μl of murine ribonuclease (RNase) inhibitor, and 100 ng of template DNA. Nuclease-free water was added to a final volume of 25 μl per reaction in 96-well plates. Reactions were incubated at 37 °C for 3 h. GFP fluorescence was measured using a BioTek Synergy H1 multimode reader (Agilent) with excitation at 480 nm and emission at 520 nm.

### Sequence-level biosecurity screening and application-specific checkpoints

The open-source IBBIS Common Mechanism^30^ (Commec v1.0.3) was used to screen nucleotide sequences in FASTA format. The complete Commec workflow comprises three functional modules: BIORISK, TAXONOMY, and LOW-CONCERN. BIORISK uses profile hidden Markov model searches implemented with hmmscan against curated biorisk profiles to identify matches associated with pathogenicity, toxin function, or virulence factors. TAXONOMY combines protein-level similarity searches using BLASTX or DIAMOND with nucleotide-level searches using BLASTN against National Center for Biotechnology Information (NCBI)-derived databases to establish taxonomic context. LOW-CONCERN uses protein, RNA, and DNA searches, implemented with hmmscan, cmscan, and BLASTN, to determine whether taxonomy-derived matches have low-concern explanations; this module does not override BIORISK matches. Native Commec outputs and annotations were retained in the screening records.

To provide access to sequence screening without dedicated high-performance computing infrastructure, Commec was deployed through a Google Colab notebook supporting environment setup, FASTA upload, screening execution, and report generation (Supplementary Fig. 3). Pre-synthesis screening of CAPE designs, pre-construction screening of GALS variants, and post-production screening for the iGEM Distribution Kit were performed using this implementation (colab.research.google.com/drive/1JvIxU67nSD5IMSvRqvCYnGl8ERaVLHCT). The notebook was also made available to CAPE participants as a no-code educational resource to promote biosecurity awareness.

Because the reference databases required for TAXONOMY exceeded the computational resources available in the Colab environment, Commec was run in taxonomy-skipping mode (--skip-tx --skip-nt). Under this configuration, BIORISK searches were performed, whereas protein- and nucleotide-level TAXONOMY searches were omitted. The absence of a BIORISK match therefore indicated only that no curated biorisk profile was matched under the screening configuration used; it did not establish the absence of taxonomic association with a regulated organism. In the study reports, records with no BIORISK match received the message “No matches found during any stage of analysis” and an overall status of *Warning*. This no-match warning was distinguished from a BIORISK *Warning* or *Flag* and was not treated as a positive BIORISK finding. The native overall status was retained and was not interpreted as comprehensive biosecurity clearance.

At each designated checkpoint, a BIORISK *Warning* or *Flag* triggered a hold and human review of the Commec annotations, sequence context, independent homology searches, and relevant reference or registry metadata. Downstream actions, including DNA synthesis, mutant construction, and kit distribution, remained on hold until review was completed and a biosecurity disposition was recorded. Incomplete or failed screening runs likewise triggered a hold until a valid report was obtained. Review-triggering records were manually inspected by authors with molecular biology expertise (Y.G. and J.Z.), and biosecurity dispositions were determined jointly with a corresponding author (T.S.). Records judged not to require restriction on biosecurity grounds could proceed subject to independent quality assurance and quality control (QA/QC) criteria and institutional requirements. Any record that remained unresolved or was judged potentially concerning would have been escalated for institutional review; no such escalation was required in this study. Record-level native outputs and human review records were compiled as Supplementary Data 2. Application-specific screening outcomes, human review findings, and downstream actions are summarized in Supplementary Table 8.

The Commec implementation was considered one component of a broader biosecurity architecture spanning laboratory access, synthesis-provider controls, and model deployment. Remotely operated automated laboratories may lower barriers to experimentation while introducing distinct misuse risks^53^; policy proposals therefore recommend customer and experiment screening, with trigger-based human review of concerning synthesis requests or protocols^54^. The 2024 Office of Science and Technology Policy (OSTP) Framework for Nucleic Acid Synthesis Screening and the International Gene Synthesis Consortium (IGSC) Harmonized Screening Protocol v3.0 address provider-side sequence and customer screening, expert review, record retention, and order escalation or refusal^59, 60^. IBBIS guidance additionally addresses verification of customer identity, legitimacy, and intended use^58^. At the model level, BioTIER (Biological Targeted Information for Exclusion and Refusal) evaluates refusal of high-risk biological requests alongside responses to benign requests^56^. These sources provide context only: this study implemented the Commec checkpoints and application-specific human review, whereas the broader safeguards described above and formal conformance with the OSTP Framework or IGSC Protocol were not assessed.

### Sequence analysis of CAPE designs in reference to FPbase proteins

To visualize the sequence space explored by the 2025 CAPE designs, dereplicated CAPE submissions (*n* = 298) and avGFP derivative sequences from FPbase (*n* = 140; retrieved on October 15, 2025) were encoded using the unmodified pretrained ESM-2 model esm2_t12_35M_UR50D^28^. Sequence embeddings were obtained by mean-pooling layer-12 residue representations, excluding special and padding tokens, to yield 480-dimensional vectors. The resulting sequence representations were combined and projected into two dimensions in a single t-distributed stochastic neighbour embedding (t-SNE)^64^ fit without prior normalization, standardization or whitening. The analysis used a perplexity of 30, principal component analysis (PCA) initialization, an automatic learning rate, 4,000 iterations, Euclidean distance and random state 42. Kernel density estimation was applied to the shared two-dimensional coordinates to visualize the distributions of the CAPE and FPbase sequence sets. The ten highest-performing CAPE variants, the avGFP parent, and the sfGFP reference were overlaid as individual points (Extended Data Fig. 4a).

In a separate database-wide analysis, amino acid sequence identity was calculated between each community design (*n* = 298 for CAPE 2025; *n* = 556 for CAPE 2026) and every protein sequence in an FPbase snapshot (*n* = 991; retrieved on August 26, 2026). Each design was globally aligned against all FPbase sequences using Needleman–Wunsch alignment with the BLOSUM62 substitution matrix and affine gap penalties of −10 for gap opening and −1 for gap extension. For each design, the closest FPbase match was defined as the sequence yielding the highest alignment score, with score ties resolved by sequence identity. Identity was calculated as the fraction of identical residues after excluding terminal-gap columns. Fig. 3g plots sequence identity to the selected match against the corresponding CFPS fluorescence fold change relative to the avGFP control measured on the same plate.

### Magnetic bead-based protein purification in microtiter plates

His-tagged proteins from CFPS reactions were purified using Ni-NTA Magnetic Beads (Smart-Lifesciences, Changzhou, China). The bead slurry was resuspended in five bead volumes of binding buffer by vortexing and dispensed into 96-well plates as 100 μl aliquots per well. Beads were equilibrated by magnetic separation on the magnetic block, removal of the supernatant, and two washes with 100 μl of lysis buffer (50 mM Tris-HCl, 500 mM NaCl, pH 8.0), with resuspension by pipetting 10 times between washes. CFPS reaction mixtures (50 μl) were added to the equilibrated beads and incubated for 30 min at room temperature with shaking at 1,000 rpm on the heater-shaker module. The beads were then magnetically separated, and the supernatant was removed. To remove nonspecifically bound proteins, the beads were washed sequentially with 150 μl of wash buffer containing 0 mM imidazole and 150 μl of wash buffer containing 20 mM imidazole. Target proteins were eluted by resuspending the beads in 150 μl of elution buffer containing 250 mM imidazole and incubating for 10 min.

### Bradford assay for protein quantification

Protein concentration was determined using the Bradford Protein Assay Kit (Beyotime Biotechnology, Shanghai, China). A standard curve was generated using bovine serum albumin (BSA) diluted in PBS to concentrations ranging from 0 to 1.5 mg ml⁻¹. For the assay, 5 μl of each standard or sample was mixed with 250 μl of G250 dye reagent in a 96-well plate. Following a 10-minute incubation at room temperature, absorbance was measured at 595 nm using a microplate reader, and protein concentrations were calculated by comparison to the BSA standard curve.

### Manual GFP expression and purification

Transformed BL21(DE3) cells were cultured in 1 litre of LB medium at 37 °C until OD_600_ reached 0.6-0.8, followed by induction with 0.5 mM IPTG at 18 °C for 16-20 h. Cells were harvested by centrifugation and resuspended in Lysis buffer (50 mM Tris-HCl, 500 mM NaCl, 10 mM imidazole, pH 8.0). Cell lysis was performed by sonication on ice, cleared lysate obtained by centrifugation (4,000 × g, 30 min, 4 °C) was applied to Ni-NTA agarose (Qiagen), and proteins were eluted with 250 mM imidazole. Following concentration using Amicon Ultra-15 centrifugal filters (30 kDa MWCO, Millipore), proteins were further purified by size exclusion chromatography on a Superdex 200 pg 16/600 column using an ÄKTA prime PLUS system (GE Healthcare) equilibrated with 25 mM Tris-HCl, 150 mM NaCl, pH 8.0, with elution monitored at 280 nm absorbance (A_280_).

### Photophysical characterization of avGFP, Var1 and sfGFP

For spectroscopic characterization of purified proteins, excitation spectra were recorded from 300-520 nm (λ_em_ = 550 nm) and emission spectra from 480-650 nm (λ_ex_ = 450 nm), both with 2 nm bandwidth. Quantum yields were determined using sfGFP as a reference standard (QY = 0.65) with integrated emission (λ_ex_ = 450 nm), normalized to absorbance at 450 nm, according to the following equation,

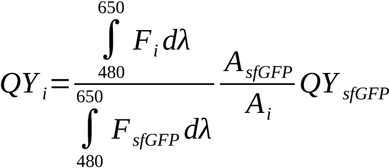

where *QY* denotes quantum yield, *F* denotes fluorescence emission intensity at wavelength λ nm, *A* denotes absorbance at 450 nm, and *i* denotes a particular GFP. Extinction Coefficient (ε) was calculated using the formula:

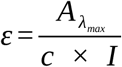

where *ε* denotes extinction coefficient, A*_λ_*_max_ is absorbance of proteins at maximum absorption wavelength, *c* is molar concentration, *I* is path length (0.7 cm). Molecular brightness was calculated as the product of extinction coefficient and quantum yield and is reported in mM⁻¹ cm⁻¹.

### Characterization of sequence-divergent CAPE 2026 designs

Nineteen sequence-divergent CAPE 2026 designs (<50% amino acid sequence identity to their closest FPbase matches) were selected for follow-up characterization because their initial CFPS signals were insufficient to establish fluorescence (Fig. 3g). Three independent *E. coli* BL21(DE3) clones per design were cultured in 24-well plates at 37 °C and 800 rpm as biological replicates, with empty-vector cultures as the negative control. Following overnight growth and subculture for 3 h, expression was induced with 0.5 mM IPTG at 18 °C and 800 rpm for 24 h. Cultures were subsequently maintained at 22 °C without shaking. Whole-cell fluorescence was monitored over time to accommodate potentially slow chromophore maturation, as reported for other fluorescent proteins generated *de novo*^65^. Cells from 100 μl culture aliquots were washed twice with PBS before fluorescence spectral scans and intensity measurements (λ_ex_ = 430 nm; λ_em_ = 510 nm); fluorescence values were not normalized to optical density. On day 4, several candidates exhibited fluorescence signals clearly above the empty-vector control (Supplementary Fig. 4a). Var2, which had the highest mean whole-cell fluorescence signal at this time point, was selected for purification as described above.

Excitation and emission spectra of purified Var2 were also recorded. Fluorescence intensities (λ_ex_ = 430 nm, λ_em_ = 510 nm) were measured for purified Var2 at 15.8 mg ml⁻¹ and avGFP at 0.1 mg ml⁻¹ alongside a buffer-only blank. Because the protein concentrations differed, these intensity measurements were used to assess fluorescence detectability rather than relative molecular brightness.

### Construction of GALS mutants

GALS variant libraries were created using a previously reported biofoundry workflow^36^, comprising 19 non-parental amino-acid substitutions at each of the two positions, giving 38 single-site (19 × 2) and 361 double-site (19 × 19) variants. First, site-directed mutations were introduced via PCR using designed primers. Following two-fragment Gibson assembly, 5 μl of reaction mix was used to transform 50 μl of competent *E. coli* DH5α in 96-well format (4 °C for 30 min, 42 °C for 60 s), which were then recovered in 200 μl of LB at 37 °C and 800 rpm for 2 h before plating on selective agar medium. Single colonies were isolated using a QPix 420 colony picker and cultured overnight in 500 μl selective LB at 37 °C and 800 rpm. For each construct, three random colonies were picked for sequence verification. Strains representing all 38 single-site and 361 double-site variants at V281 and F282 were then selected by hit picking. Plasmid DNA was extracted on a KingFisher Flex module using the 96 Mag-MK Plasmid DNA Mini-Preps Kit (TIANGEN, Beijing, China). To create strain libraries, plasmid DNA was then used to transform BL21(DE3)-Δ6, and two independent clones were selected for subsequent phenotypic screening.

### Whole-cell FALD biotransformation for mutant screening

Mutant colonies were picked into 96 deep-well microplates (Jetbiofil, Guangzhou, China) containing 200 μl of LB+Kan and incubated for 24 h at 37 °C. Cultures were diluted 1:100 into 1 ml of fresh LB+Kan containing 0.1 mM isopropyl β-D-1-thiogalactopyranoside (IPTG) and incubated at 37 °C for 4 h followed by 20 h at 30 °C. Cells were harvested by centrifugation at 3,300 × *g* for 5 min, washed twice using phosphate-buffered saline (50 mM PBS, pH 7.4), and resuspended in assay buffer (50 mM PBS, 5 mM MgSO**_4_**, 0.5 mM thiamine pyrophosphate (TPP), 50 mM (1.5 g l⁻¹) FALD, pH 7.4). After incubation at 30 °C for 24 h, supernatants were collected after centrifugation at 3,300 × *g* for 5 min for product analysis.

### Spectrophotometric assays for GALD and DHA quantification

For spectrophotometric GALD detection^36^, diphenylamine (DPA) reagent was prepared by dissolving 1.5 g DPA in 100 ml of glacial acetic acid followed by addition of 1.5 ml of concentrated sulfuric acid. Sample and standard solutions (30 μl) were mixed with 150 μl of DPA reagent in 96-well PCR plates (Jetkeen, Dongguan, China), incubated at 90 °C for 30 min, and transferred to 96-well microtiter plates (Corning 3599) for endpoint absorbance measurement at 650 nm. The assay was linear over a GALD concentration range of 3.125 to 200 mg l⁻¹ (0.052 to 3.333 mM). For spectrophotometric DHA detection^36^, 90 μl of sample or standard solution was dried at room temperature in 96-well microtiter plates in a chemical fume hood. Samples were reconstituted in 90 μl of 50 mM PBS (pH 7.4), followed by addition of 60 μl Buffer A (0.2 mg ml⁻¹ galactose oxidase, 24 U ml⁻¹ horseradish peroxidase, 5 mM MgSO_4_ in 50 mM PBS, pH 7.4) and then 50 μl Buffer B (3.2 mM ABTS, pH 7.4). Absorbance at 410 nm was recorded every 30 s for 20 min, and the initial linear slope (ΔA_410_ min⁻¹) was used for quantification. The assay was linear over a DHA concentration range of 0.0625 to 25 mg l⁻¹ (0.694 to 277.5 μM). Samples were diluted as necessary to fall within the validated linear range of each assay, and measured concentrations were corrected for the dilution factor.

### Manual GALS Protein Expression and Purification

Genes encoding the parent and mutant GALS were assembled into the pET28m expression vector by Gibson Assembly. *E. coli* BL21(DE3) cells carrying recombinant plasmids were cultivated in 5 ml of LB+Kan at 37 °C for 16-20 h and subsequently used to inoculate 1-litre cultures. When OD_600_ reached 0.6-0.8, IPTG was added to a final concentration of 0.5 mM, followed by 18 °C induction for 20 h. Cells were harvested by centrifugation at 8,000 × *g* and resuspended in 50 ml of lysis buffer (500 mM NaCl, 50 mM Tris-HCl, 5 mM MgSO_4_, 10% (v/v) glycerol, pH 7.5).

Cell lysis was performed using a high-pressure homogenizer (JNBIO, Guangzhou, China), and debris was removed by centrifugation at 10,000 × *g* for 30 min at 4 °C. The clarified lysate was loaded onto a nickel affinity column (GE Healthcare), washed with 50 ml of wash buffer (150 mM NaCl, 25 mM Tris-HCl buffer, 5 mM MgSO_4_, and 20 mM imidazole, pH 7.5) and eluted with 20 ml of elution buffer (150 mM NaCl, 25 mM Tris-HCl, 5 mM MgSO_4_, and 250 mM imidazole, pH 7.5). Purified protein was concentrated and buffer-exchanged into 50 mM PBS with 5 mM MgSO_4_ (pH 7.4) using an Amicon Ultra centrifugal filter device (Millipore, USA, 30 kDa MWCO). Protein concentration was determined using the Bradford Protein Assay Kit (Beyotime).

### *In vitro* Kinetic Characterization of Purified GALS Variants

Reaction mixtures (200 μl) contained 1 mg ml⁻¹ purified GALS, 66.7 mM (2.0 g l⁻¹) FALD, 5 mM MgSO_4_ and 1 mM TPP in 50 mM PBS (pH 7.4) and were incubated at 37 °C for 3 h. Reactions were terminated by heating at 80 °C for 10 min. After centrifugation at 4,000 × *g* for 5 min, the supernatants were collected and GALD and DHA were quantified using the spectrophotometric assays described above. FALD conversion was calculated from product formation according to reaction stoichiometry as 100% × (2[GALD]+3[DHA])/[FALD]_0_. Three independent reactions were performed for each variant.

To validate the hits identified in the whole-cell biotransformation screen, purified GALS variants were assayed for the enzymatic conversion of FALD to GALD and DHA. Kinetic parameters were determined for parent GALS and the W/L (V281W/F282L) variant in 200 μl reactions performed in 96-well microplates. Initial rates of GALD and DHA formation were measured at 37 °C in 50 mM PBS (pH 7.4) containing 5 mM MgSO_4_ and 0.5 mM TPP upon addition of 1 mg ml⁻¹ purified enzyme and expressed as FALD-equivalent rates based on reaction stoichiometry (2*v*_GALD_ and 3*v*_DHA_, respectively). FALD concentrations ranged from 20 to 1,500 mM (0.6 to 45.0 g l⁻¹). Reactions were terminated after 20 min by heating at 80 °C for 10 min. After centrifugation at 4,000 × g for 5 min, the supernatants were collected, and GALD and DHA were quantified using the spectrophotometric assays described above. For estimation of apparent kinetic parameters^36^, only data within the pre-inhibition range were used for fitting. GALD-derived rates were fitted to the Michaelis–Menten model, whereas the DHA-derived rates were fitted to the sigmoidal Hill equation, using GraphPad Prism 9 software to process the pre-inhibition range data. The combined FALD-equivalent rate was calculated as *v*_total_ = 2*v*_GALD_ + 3*v*_DHA_ (Fig. 4d). Three independent reactions were analyzed at each enzyme-substrate concentration combination.

### Two-stage Cell-free Enzyme Cascade for EG Production

A decoupled two-stage enzyme cascade^37^ was used to convert FALD to EG. First-stage reaction mixtures (200 μl) contained 50 mM PBS (pH 7.4), 333 mM FALD (10 g l⁻¹), 2 mM MgSO₄·7H₂O, 1 mM TPP, 1 mM dithiothreitol (DTT), and either parent GALS or the W/L variant at a final concentration of 9 mg ml⁻¹. Reactions were incubated at 37 °C for 4 h. In the second stage, GOX0313, an NADH-dependent alcohol dehydrogenase from *Gluconobacter oxydans*^37^, and NADH were added to final concentrations of 1 mg ml⁻¹ and 70 mM, respectively. The GALD generated by GALS was reduced to EG during a further 8-h incubation, whereas GOX0313 was omitted from the control reactions. GOX0313 was recombinantly expressed and purified using the procedure described above for GALS.

Reactions were quenched by adding an equal volume of 0.2 M H₂SO₄ and clarified by centrifugation at 21,130 × *g* for 10 min. The supernatants were transferred to chromatography vials and analyzed by high-performance liquid chromatography (HPLC) using an Agilent 1260 system equipped with a refractive index detector (RID). An Aminex HPX-87H column (300 × 7.8 mm; Bio-Rad) was maintained at 65 °C and eluted isocratically with 5 mM H₂SO₄ at a flow rate of 0.6 ml min⁻¹. The injection volume was 5 μl. GALD, FALD, EG, and formic acid (FA) were resolved in the same chromatographic run and identified by comparing their retention times with those of authentic standards. GALD and EG were quantified using RID, with compound-specific external calibration curves spanning 0.01–5.0 g l⁻¹. Three independent cascade reactions were analyzed for each GALS enzyme.

### Automated Production and Quality Assurance of the iGEM Distribution Kit

Dried plasmid DNA samples were recovered through *E. coli* transformation, colony isolation, overnight cultivation, and plasmid extraction using the previously reported biofoundry workflow^36^ described above for GALS library construction. Frozen strain stocks were prepared by mixing 100 μl of cell culture with 100 μl of 40% glycerol and storing the mixtures at −80 °C. Plasmid DNA concentrations were measured using a Stunner instrument (Unchained Labs, Pleasanton, CA, USA). DNA samples were normalized to 1 ng μl⁻¹ using 0.1% cresol red in TE buffer in 96-deep-well master plates. Aliquots of 3 μl were dispensed into 384-well destination plates and air-dried, after which the plates were foil-sealed before shipment.

Whole-plasmid sequencing data were analyzed using a custom Biopython pipeline for global alignment against reference sequences, mismatch enumeration, and structural variant detection. Ambiguous alignments were manually inspected in SnapGene v6.0.2. All variant-containing and no-alignment records were subsequently screened using the Commec Colab implementation described above. Records documenting iGEM clone identity, manufacturing QA/QC, and kit-release decisions were compiled as Supplementary Data 4.

